# The Mechanism and Energetics of the Dynein Priming Stroke

**DOI:** 10.1101/2023.06.10.544469

**Authors:** Mert Golcuk, Sema Zeynep Yilmaz, Ahmet Yildiz, Mert Gur

## Abstract

Dyneins is an AAA+ motor responsible for motility and force generation towards the minus end of microtubules. Dynein motility is powered by nucleotide-dependent transitions of its linker domain, which transitions between straight (post-powerstroke) and bent (pre-powerstroke) conformations. To understand the dynamics and energetics of the linker, we per-formed all-atom molecular dynamics (MD) simulations of human dynein-2 primed for its power stroke. Simulations re-vealed that the linker can adopt either a bent conformation or a semi-bent conformation, separated by a 5.7 kT energy bar-rier. The linker cannot switch back to its straight conformation in the pre-powerstroke state due to a steric clash with the AAA+ ring. Simulations also showed that an isolated linker has a free energy minimum near the semi-bent conformation in the absence of the AAA+ ring, indicating that the linker stores mechanical energy as it bends and releases this energy during the powerstroke.

## INTRODUCTION

Dyneins are large (1.4 MDa) motor proteins that perform most of the minus-end directed motility and force generation functions on microtubules (MTs). Cytoplasmic dynein (dynein hereafter) transports a wide variety of intracellular cargos and performs essential functions during cell division. Other dyneins are localized to cilia where they drive intraflagellar transport in the retrograde direction and power ciliary beating.^1^ The partial loss of dynein function has been associated with numerous developmental and neurodegenerative diseases and ciliopathies.^2,3^

Dynein motility is driven by the C-terminal motor domain of the dynein heavy chain.^2,4,5^ The motor domain is comprised of six AAA+ modules (AAA1-6) that are arranged into a hexagonal ring.^6^ AAA1-4 modules contain conserved the Walker A and B motifs required for ATP binding and hydrolysis, whereas AAA5-6 plays a structural role.^7,8^ Catalytic activity of AAA1 is essential for dynein motility,^9,10^ while nucleotide binding and hydrolysis at AAA2-4 were proposed to have structural and regulatory roles.^9,11^ The AAA+ ring connects to a small globular MT-binding domain (MTBD) through a 15 nm long antiparallel coiled-coil stalk that extends between AAA4 and AAA5.^6^ The stalk is supported by another coiled-coil, named buttress,^12^ which facilitates communication between the AAA+ ring and MTBD by controlling the registry of stalk coiled-coils.^4,13^ The surface of the AAA+ ring form multiple contacts with an α-helical bundle, named linker. The linker can be divided into two rigid subdomains connected by a flexible hinge (H10).^7,14^ While Link1-2 (helices H5-H9) undergoes rigid body motion throughout the dynein mechanochemical cycle, Link3-4 (helices H11-H18) stays rigid and in contact with AAA1 (Fig. 1a).^15^

**Figure 1.**
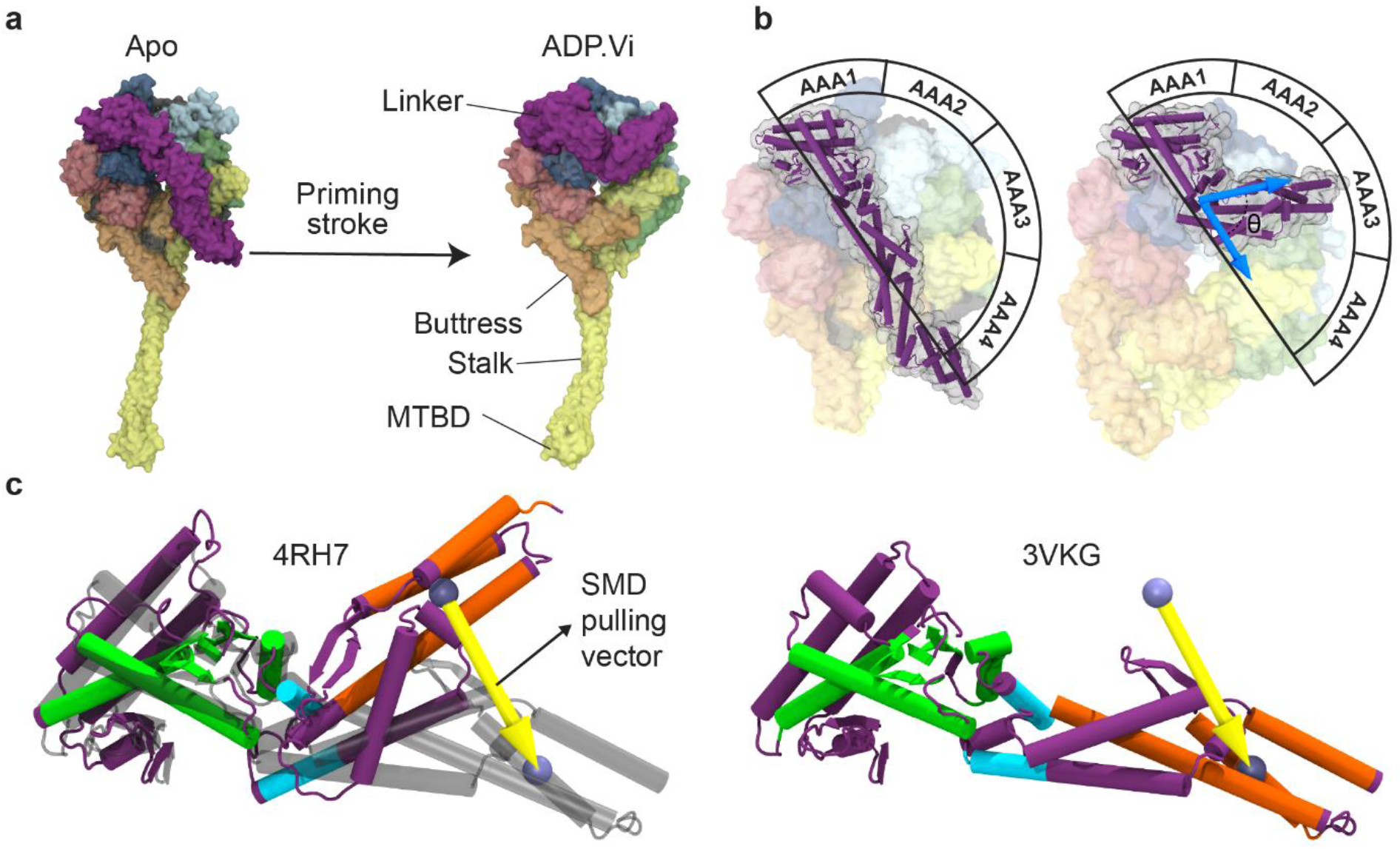
Nucleotide-dependent conformational changes of the linker domain. **a**, Crystal structure of the cytoplasmic dynein motor domain in the ADP state with straight linker (PDB ID: 3VKG^7^, left) and in the ADP.Vi state with bent linker (PDB ID: 4RH7^15^, right) shown in surface representation. **b**, The angle of the bent and straight linker on the AAA+ ring surface. Alpha helices (H4-H18) and beta sheets (S1-S8) of the linker are highlighted in the cartoon representation. **c**, The bent linker of the ADP.Pi-bound human dynein-2 structure superimposed onto the straight linker of the ADP-bound *D. discoideum* dynein via its Link3-4 helices. Regions comprising SMD, fixed atoms for SMD simulations, and the atoms used in the alignment of the linker are shown in orange, turquoise, and green, respectively. The center of mass (CoM) of SMD atoms in bent and straight linker structures are shown with ice and blue spheres, respectively. SMD pulling vector pointing from the CoM of SMD atoms in the bent linker to that in the straight form is shown as a yellow arrow.

Several conformational states of the motor domain have been solved at near-atomic resolution: *Saccharomyces cerevisiae* dynein-1 in the nucleotide-free (apo) state (PDB ID: 4AKG^14^), human cytoplasmic dynein-2 in the presence of ADP.Vi, the ATP analog that mimics the ATP-hydrolysis transition state (ADP.Pi; PDB ID: 4RH7^15^), and *Dictyostelium discoideum* dynein-1 in the ADP-bound state (PDB ID: 3VKG^7^). Based on these structures and bulk FRET studies, the proposed mechanochemical cycle of dynein involves coordination between the nucleotide state of AAA1, registry of the stalk coiled coils, and conformation of the linker (Fig. S1). In the apo state of AAA1, the stalk is in an α-registry where its MTBD is strongly bound to the MT, the catalytic ring is in the open conformation, and the linker is straight, exiting the AAA+ ring at AAA4/AAA5.^7,14^ ATP binding to AAA1 closes the gap between AAA1 and AAA2, and the AAA+ ring adopts its closed conformation by upward movement of AAA4 towards the linker, and movement of AAA5 and AAA6 towards each other.^11^ Closing of the AAA+ ring triggers the release of MTBD from MT^16^ by shifting the stalk coiled-coils to the β registry^17^ and movement of the linker to its bent conformation, exiting the ring at AAA2.^1,15^ Bending of the linker is referred to as the priming stroke, which generates the net bias in dynein stepping towards the minus-end.^7,14,18^ After ATP hydrolysis, dynein rebinds the MT, which may shift stalk coiled-coils back to the α registry and trigger the release of inorganic phosphate from AAA1. Upon phosphate release, the AAA+ ring switches back to the open conformation and the linker moves from bent to straight conformation, docking near AAA4.^1^ This transition is referred to as powerstroke,^19^ which produces mechanical work required for dynein to pull its cargo in the forward direction.^20^ After ADP release, the linker moves into a slightly more extended straight conformation, thus resetting the mechanochemical cycle.^11^

The exact order of events for the priming and power stroke remains unclear, mainly due to inherent challenges in trapping AAA1 in the ATP-bound state and the presence of other ATP-binding sites in the AAA+ ring.^1^ It also remains unclear whether the linker can store elastic strain energy in its bent conformation that could be released during the powerstroke. All-atom MD simulations are ideally suited to investigate the full range of dynein conformations sampled along the mechanochemical cycle, closing the gap between high-resolution snapshots of dynein obtained by cryo-electron microscopy (cryo-EM) and conformational dynamics of dynein obtained by single-molecule imaging at lower resolution. Previously, Kamiya et al.^21^ performed a total of 200 ns all-atom MD simulations to show that the stalk is more flexible in the ATP-bound state of AAA1 relative to the ADP-bound state. More recently, we performed^22^ 1 µs long all-atom MD simulations of native and mutant dynein constructs to engineer a plus-end directed dynein. However, all-atom MD simulations could not be used to investigate the conformational dynamics of dynein due to its complexity and large size, most computational studies of dynein were limited to coarse-grain models.^23-27^

In this study, we explored the conformational dynamics and energetics of the priming stroke of the linker by performing a total of ∼16.5 µs of all-atom conventional MD, steered MD^28^ (SMD, Fig. 1c), and umbrella sampling MD^29^ (UMD) simulations of the motor domain of human dynein-2 having its AAA+ ring in the closed conformation (800k atoms in size). We also performed 9 µs of SMD and UMD simulations of the isolated linker (300k atoms in size) in the presence of explicit water and ions. These simulations revealed two free energy minimums for the linker: a semi-bent linker and a bent linker, which have similar energy levels but are separated by an energy barrier. The linker is sterically excluded to adopt its straight conformation at the surface of the closed AAA+ ring, underscoring the critical role of rigid body motions of the AAA+ ring to coordinate the conformational changes of the linker. In the absence of the AAA+ ring, an isolated linker energetically prefers a conformation close to the newly discovered semi-bent conformation, and its energy increases more than 3 *kT* when it transforms to the bent conformation, suggesting that the linker stores elastic strain energy during its priming stroke.

## RESULTS

### The linker is sterically excluded by the AAA+ ring to adopt a straight conformation

Because the most complete dynein structure, while conducting this study, was captured in the ADP.Pi state of AAA1,^15^ we mapped the conformational transition of the priming stroke by starting from this conformation and setting the straight conformation of the linker in the ADP state.^7^ We performed consecutive cycles of SMD simulations followed by constrained MD (cMD) simulations to cover the ∼46.2 ° clockwise rotation of Link1-2 relative to Link3-4 as the linker moves from its bent to straight conformation at the surface of the AAA+ ring having a closed conformation (Fig. 1b). In each SMD simulation, the linker was pulled from its N-terminal tip towards its straight structure (PDB ID: 3VKG^7^) with a 1 Å/ns speed for a duration of 5 ns, while allowing it to move freely in orthogonal directions. After each SMD, the linker was constrained for 5 ns to allow the remaining dynein structure to optimize in this new linker conformation (Fig. S2). The SMD pulling vector was updated before each SMD simulation.

After 8 consecutive cycles of SMD and subsequent MD simulations, the root mean square deviation (RMSD) of the linker from its straight conformation decreased from 9.7 Å to 3.2 Å (Fig. S1). Additional cycles were not able to further decrease the RMSD due to a steric clash between the linker and the AAA+ ring, which stayed in its closed conformation throughout simulation cycles (Fig. 2). The steric clash comprised contacts between H5, H6, H7, and H10 of the linker and the PS-I insert of AAA4 (Fig. 2 and Fig. S3-4), as predicted by structural studies.^15^

**Figure 2.**
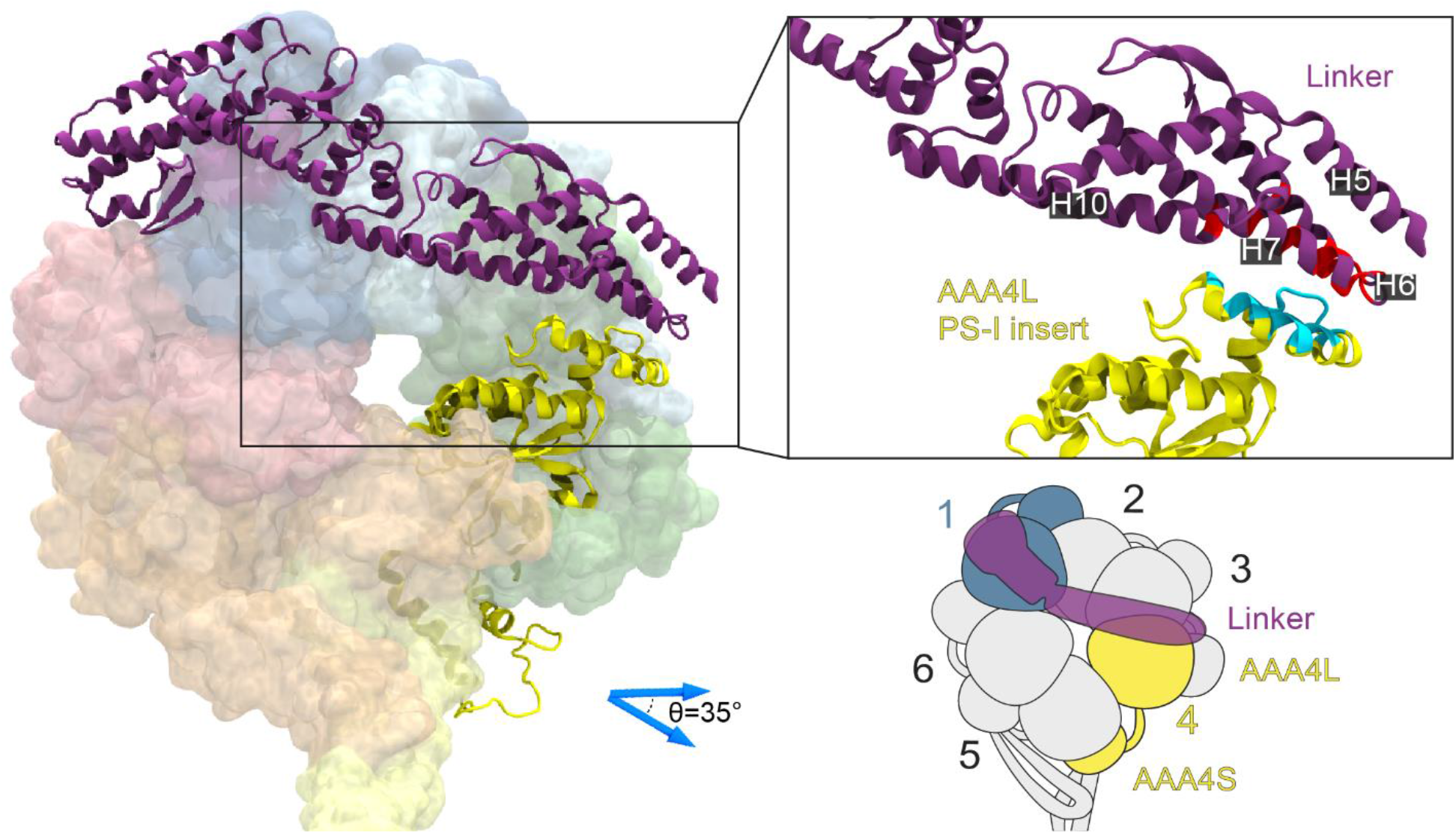
Steric clash between the linker and AAA+ ring observed in SMD simulations. The left panel shows the AAA+ ring of the dynein obtained conformations after 80 ns of SMD-cMD simulations. Linker and AAA4 are shown in cartoon representation whereas the remaining structural components are shown in surface representation. The right top panel shows the steric clash between linker residues (red): R1306, Q1310, K1313, D1314 (H5); P1316, Y1318, K1319 (H6); S1315 (loop connecting H5-H6); E1322, DP1323, V1325, S1326, E1329, R1330, A1333, E1337 (H7); R1398(H10); and AAA4L PS-I insert residues (cyan) E2742, E2745, P2746, L2748, L2749, P2750, K2752, D2753, S2756, F2761, G2762, P2763, V2764, and F2765. The right lower panel shows the schematic representation of the linker position when a steric clash was observed.

### The linker adopts two conformational states at the surface of a closed AAA+ ring

We next ran a large set of UMD simulations, totaling 5.25 μs in length, to optimize the modeled priming stroke transition and explore its energetics. RMSD of the linker to the straight conformation was selected as the reaction coordinate, ξ (see Methods). 15 UMD simulations were initiated, each restraining the linker at different RMSD values, ranging from 3.0 to 10.8 Å along the reaction coordinate, via a harmonic potential. The reaction coordinate was then subdivided into 0.1 Å wide windows and conformations sampled in UMD simulations were clustered into these windows based on their RMSD values. Using the weighted histogram analysis method (WHAM),^30^ the free energy surface for the linker during the priming stroke was constructed as a function of RMSD (Fig. 3a and Fig. S5) and projected on the angle (θ) between Link1-2 and Link3-4 (Figs. 1b and 3b). Simulations revealed two energy minima for the linker on the closed AAA+ ring: The semi-bent conformation exits the ring near AAA3/AAA4 at ξ = 4.25 Å and θ = 37° and the bent conformation exits the ring near AAA2/AAA3 at ξ = 9.25 Å and θ = 78.5° (Fig. 3c). These minima have nearly identical free energies, differing only by 0.5 *kT*, and transitions between these two conformations are limited by an energy barrier of 5.7 *kT* at θ = 63.3°.

**Figure 3.**
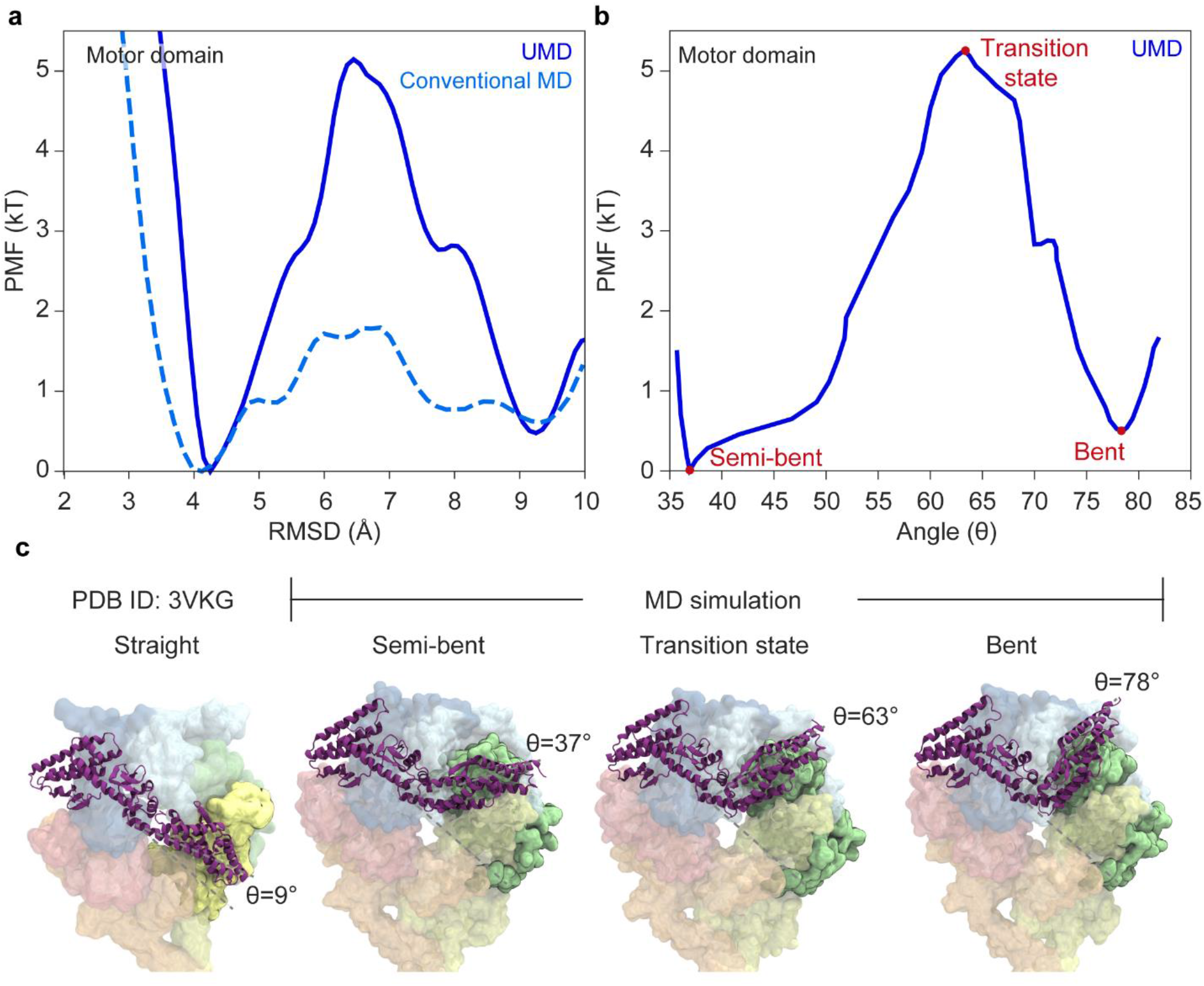
Free energy profiles of the linker priming movement. **a**, Free energy surfaces of the linker priming movement were obtained as a function of the RMSD to the bent linker structure^7^. RMSD = 0 Ǻ corresponds to the straight conformation of the linker in the post-powerstroke state. **b**, Free energy surface projected onto the angular orientation of the linker. **c**, Priming stroke of the linker based on structures and MD simulations. Illustrative representation of the priming stroke along the reaction coordinate. From left to right: Open AAA+ ring with a straight linker (PDB ID: 3VKG^7^), closed AAA+ ring with semi-bent conformation (energy minimum), transition state of the linker (energy barrier), and bent linker (energy minimum). The closed AAA+ ring conformations were sampled from UMD simulations.

Movement of the linker from the semi-bent to the bent conformation involves rearrangement of interdomain (linker-AAA+ ring) and intradomain (linker-linker) interactions (Fig. 4, S6, and Table S1-4). As the linker moves from the semi-bent conformation to the transition state, a total of five salt bridges, six electrostatic interactions (including hydrogen bonds), and six hydrophobic interactions that were observed at high frequency in the semi-bent conformation were either completely lost or their observation frequency decreased substantially. The passage through the energy barrier comprises the formation of seven new salt bridges, four electrostatic and six hydrophobic interactions, all of which were neither observed in the semi-bent nor bent conformation. Among these temporary interactions, four salt bridges, two electrostatic, and four hydrophobic interactions are only observed after the linker passes through the energy barrier in the priming stroke direction. After passing through the barrier, the linker formed twelve new salt bridges, six electrostatic and six hydrophobic interactions in the bent form, and the remaining interactions particular to the bent state were formed in later stages in simulations. We noticed that the number of high-frequency interactions was lower in the transition state than that observed in semi-bent and bent conformations, providing a possible explanation for the free energy barrier between these conformations.

**Figure 4.**
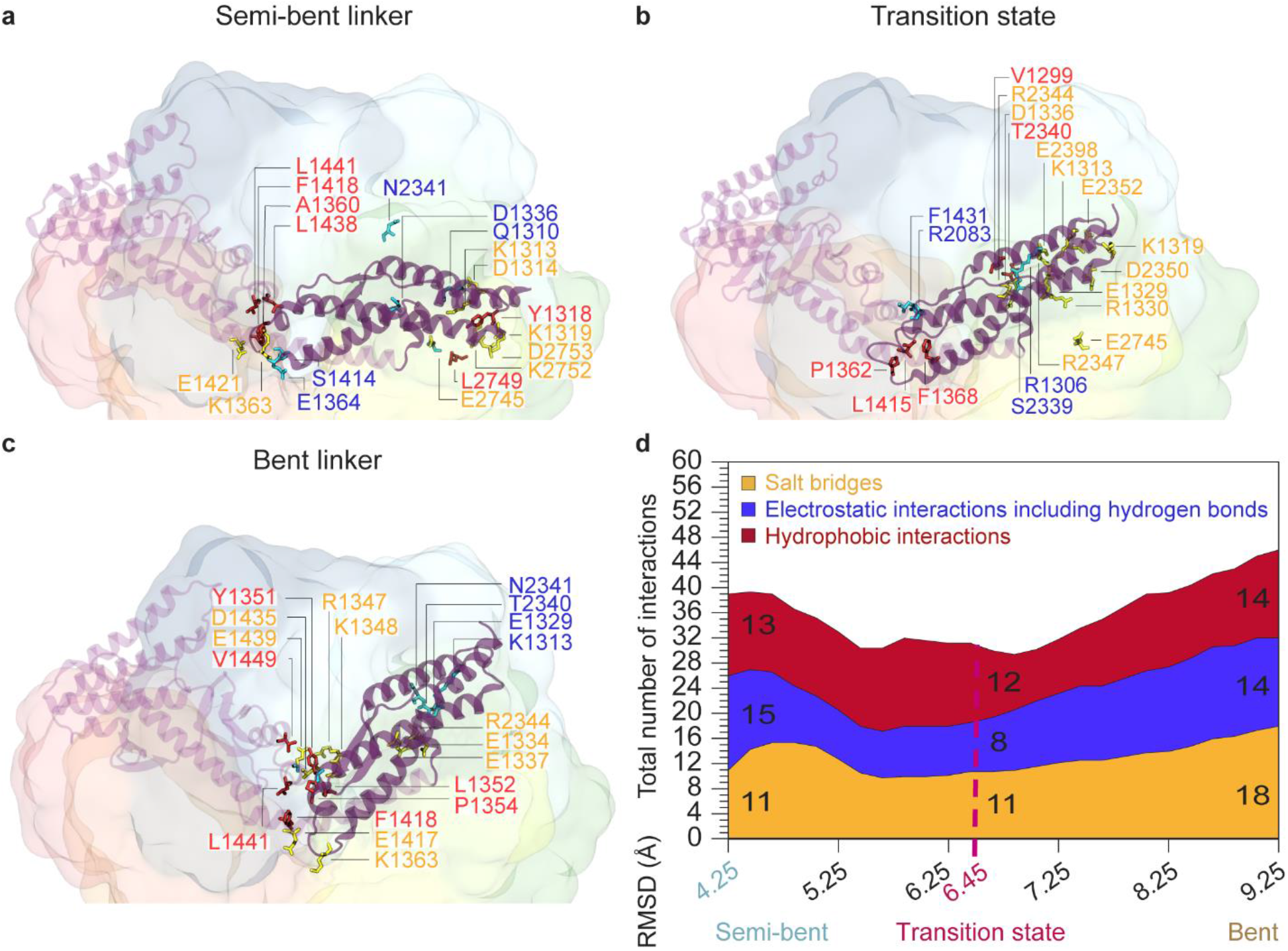
Change in the total number of intra- and inter-domain interactions of the linker during its priming movement. **a-c**, High-frequency pairwise interactions were observed for the semi-bent, transition state, and bent conformations of the linker. Only the unique interactions that are observed for a given conformation are shown. See Fig. S6 for medium-frequency interactions. **d**, Changes in the total number of salt bridges, electrostatic interactions (including hydrogen bonds), and hydrophobic interactions observed with high frequency, due to the formation and breakage of these interactions during the priming stroke. See Fig. S7 for a breakdown of all interactions into intra- and inter-domain interactions, including medium-frequency interactions, and Fig. S8 for the change in the total number of interactions based on the linker angle.

### Bending of an isolated linker is an energetically uphill process

We next asked whether the linker can store elastic strain energy as it bends during the priming stroke. We isolated the bent conformation of the linker from the motor domain conformations sampled in our MD simulations. The transition of the linker between its bent to straight conformation was modeled by biasing the bent linker conformation towards its straight “target” conformation via targeted MD (TMD) simulations in the absence of the AAA+ ring (see Methods).^31^ To construct the free energy surface of the isolated linker, we performed a total of 9 µs ns long UMD simulations^29^ using conformations sampled from TMD simulations, each separated by 0.5 Å in RMSD.

The location of free energy minima conformation of an isolated linker (ξ=3.5 Å, Fig. 5 and Fig. S5) was similar to the semi-bent conformation we observed in the presence of the AAA+ ring. Thus, the isolated linker does not energetically favor the bent conformation observed in the pre-powerstroke state. We only observed a shallow minimum that overlapped with the location of the bent conformation, but there was a large (3.3 *kT*) free energy difference between the energy minima of semi-bent and bent conformations. Thus, bending of the isolated linker is an energetically unfavorable process when its surface is in complete contact with explicit water.

**Figure 5.**
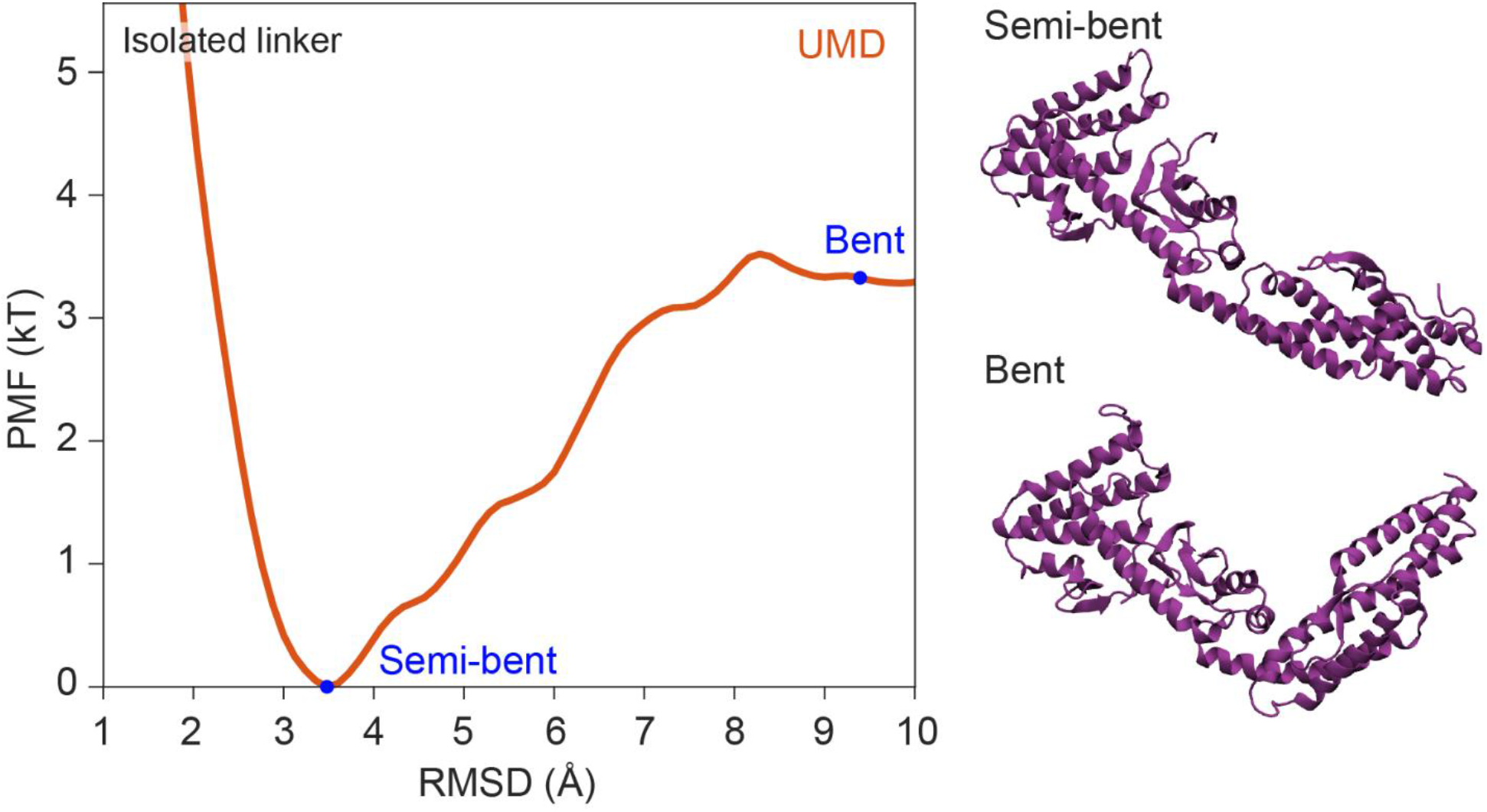
Free energy profile of the isolated linker. The free energy profile of an isolated linker along the reaction coordinate (left) was constructed based on UMD simulations of the linker performed in the absence of the AAA+ ring (right).

For the isolated linker, the net change in the number of high-frequency interactions was +2 salt bridges, 0 electrostatic, and -1 hydrophobic interaction when the linker was moved from the semi-bent (ξ=3.5 Å) to bent (ξ=9.3 Å) conformation (Fig. S7). In comparison, in the presence of the AAA+ ring, movement of the linker from the semi-bent (ξ=4.3 Å) to bent (ξ=9.3 Å) conformation exhibited a change of +7 salt bridges, -1 electrostatic, and +1 hydrophobic interaction. Because the linker has a larger net change of favorable interactions in the presence of the AAA+ ring, we concluded that the bent conformation of the linker in the pre-powerstroke state is stabilized by its interactions with the close conformation of the AAA+ ring.

### Unconstrained MD simulations initiated from intermediate linker conformations

To validate the energy surface generated via UMD simulations and sample the conformational space in the absence of external bias, we performed 75 unbiased conventional MD simulations, each of 150 ns length, initiated from UMD simulation conformations that are equally separated by 0.1 Å along the reaction coordinate. Although linker movements were observed to varying degrees in the simulations, none of them fully captured the complete transition of the linker from its straight to bent conformation. We then combined these simulation trajectories into a single 11.25 µs long MD trajectory. Sampled conformations were clustered based on their linker RMSDs to the straight conformation. Since distributions were obtained without an external biasing potential, a direct approach^32^ was used to estimate the free energy surface (Fig. 6, Methods). The locations of the free energy minima for the semi-bent (ξ=4.1 Å) and bent (ξ=9.4 Å) conformations, and the transition state (ξ=6.7 Å), as well as the free energy difference (0.61 *kT*) between the semi-bent and bent conformations, were nearly identical to those obtained from UMD simulations (Fig. 3a). The free energy barrier (1.8 *kT*) was lower than that obtained from UMD simulations. This is presumably because we initiated 21 unbiased conventional MD simulations in the vicinity of the transition state (ξ=6.5-7.5 Å), which is an energetically unfavorable low probability region (Fig. 3a) and would otherwise be sampled with considerably lower frequency. The simulations initiated from the proximity of the energy barrier quickly moved towards the closest minimum (Fig. S9).

**Figure 6.**
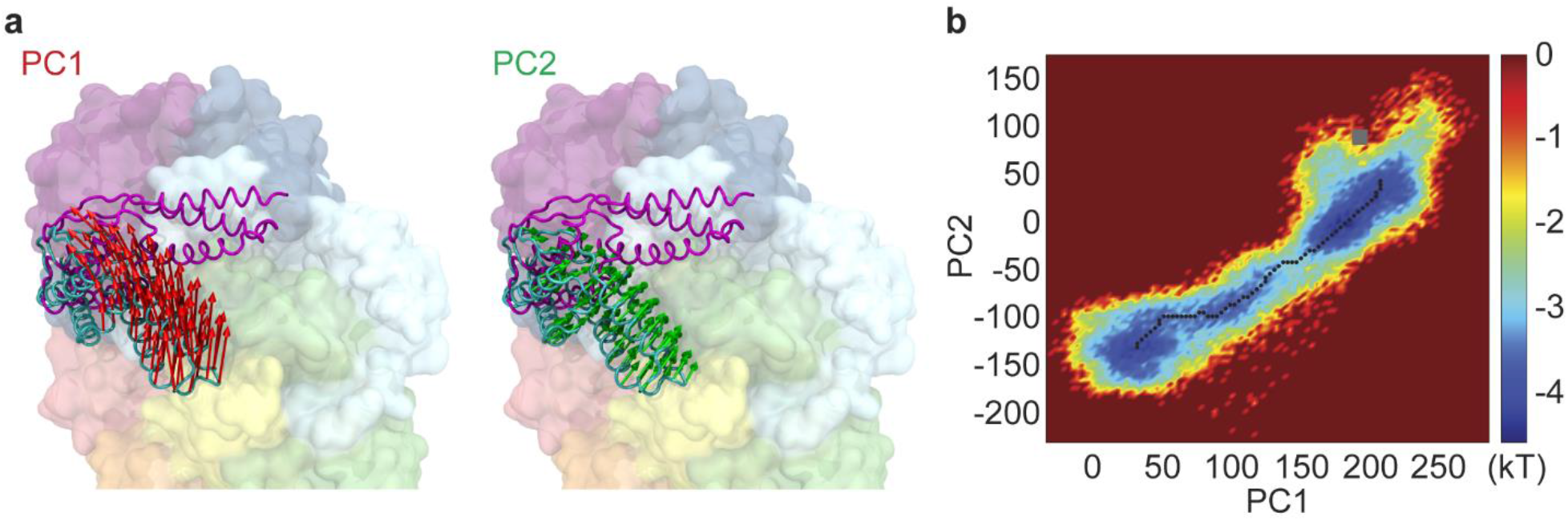
Two most prominent linker movements and the projected free energy landscape obtained from unbiased MD simulations. **a**, The directions of movement corresponding to PC1 (red) and PC2 (green), obtained from conventional MD simulations, are indicated by arrows shown on the bent linker conformation (grey). The straight conformation of the linker is superimposed via its Link3-4 subdomains. **b**, The free energy landscape constructed from cMD simulations. The reaction coordinates describing the x and y axes were defined as PC1 and PC2, respectively. The minimum free energy pathway connecting the bent and straight linker states is shown with dots.

We performed the first principal component analysis (PCA)^33^ to obtain distributions along the most prominent motions of the linker in the combined 11.25 µs trajectory of unbiased MD simulations (see Supplementary Information). Subsequently, conformations were projected onto the first two principal components (PCs), PC1 and PC2 (Fig. 6a), which cumulatively accounted for 95.2% (72.1% for PC1 and 23.1% for PC2) of rigid body motions of Link1-2 (Fig. 6b). Based on these projections, we constructed the minimum free energy pathway for the transition of the linker from the semi-bent to the bent conformation (Supplementary video 1), which we propose as the transition pathway of the motor domain from its semi-bent linker form to its bent linker form.

## DISCUSSION

In this study, we investigated the mechanism and energetics of the linker movement between its bent and straight conformations by performing all-atom MD simulations. UMD simulations revealed two energy minima, which correspond to bent and semi-bent conformations of the linker on the surface of a closed conformation of the AAA+ ring. The linker docks onto AAA2/3 in the bent conformation and onto AAA3/4 on the semi-bent conformation we observed. The free energy surface obtained via UMD and conventional MD simulations revealed that the transition from the semi-bent to bent conformation, covering more than half of the priming stroke (54% in terms of RMSD and 53% in terms of linker angle), is an energetically neutral process. Therefore, the linker can adopt either of these two conformations in the pre-powerstroke state, but transitioning between these conformations is limited by an energy barrier of 5.7 *kT*. This conformational heterogeneity of the linker in the pre-powerstroke state is consistent with prior cryo-EM studies, which either could not clearly resolve the linker density^34^ or observed the linker to dock onto either AAA2/3 (bent) or AAA4 (similar to the semi-bent conformation)^11^ and also with coarse-grained simulations of dynein with implicit nucleotides that showed the linker of ATP-bound dynein to spend a significant portion of time in a partially bent state during its priming stroke.^23^

UMD simulations demonstrated that the bending of a linker is an energetically uphill process, suggesting that the linker is a mechanical element that stores energy internally during the priming stroke. These results are consistent with a negative stain EM study, which showed that the linker decoupled from the surface of the AAA+ ring of *D. discoideum* dynein prefers to adopt a near-straight conformation, whereas bent conformations were not observed.^34^ Despite this conformational preference, semi-bent and bent conformations of the linker in the pre-powerstroke state are stabilized with an increased number of salt bridges and hydrophobic interactions between the linker and the closed conformation of the AAA+ ring. These results indicate that linker-AAA+ ring interactions contribute to the energetics of the dynein mechanochemical cycle by allowing the linker to store mechanical energy in its pre-powerstroke conformation. We also observed that an isolated linker does not prefer to switch to a fully straight post-powerstroke conformation, indicating that transition of the linker from the bent to semi-bent conformation releases mechanical energy, but the transition from the semi-bent to fully-straight conformation is an energetically uphill process stabilized by the interactions with the linker and the AAA+ ring.

Steering the bent linker toward its straight conformation on the surface of the closed conformation of the AAA+ ring showed that the linker is not able to reach the fully extended post-powerstroke conformation due to a steric clash between the linker and the AAA4 site. These results are consistent with the proposed steric clash between the linker of the ADP-bound dynein^7^ and the AAA4 module of the ADP.Vi-bound dynein.^15^ The alignment of these two structures with respect to the static Link3-4 part of the linker (Fig. S3) showed that the PS-I insertion of AAA4 serves as a mechanical gate that prevents the powerstroke of the linker before the ring adopts its open conformation through rigid body arrangement of its AAA+ modules (Fig. S4).^15^ Similarly, superimposing the linker of ADP.Vi-bound dynein with the AAA+ ring of the ADP-bound dynein results in a steric clash with the bent linker and the AAA3 module of the AAA+ ring (Fig. S3), indicating that AAA3 serves a gate that prevents the priming stroke of the linker until ATP binds to AAA1 and closes the AAA+ ring. Thus, the transition of the AAA+ ring from its closed to open conformation causes the AAA4 site to push the linker towards AAA3 while simultaneously lifting the AAA3 gate for the priming movement of the linker towards AAA2. Opening of the AAA+ ring also prevents the linker’s reverse movement to the straight conformation via a steric clash at the AAA4 site, which could be critical for maintaining a productive mechanochemical cycle. It remains to be determined whether the open conformation of the AAA+ ring establishes a new energy minimum surface that stabilizes the linker in its straight conformation during the powerstroke.

## METHODS

### System Preparations and Details of MD simulations

Three sets of MD simulations were performed starting from the dynein structure captured in the ADP.Pi state of AAA1.^22^ The study^22^ selected the human dynein-2 motor domain structure primed for power stroke (PDB ID: 4RH7^15^) as starting structure. The structure was solvated in a water box using the TIP3P water model, having at least 15 Å of water padding in each direction (i.e., protein has at least 30 Å distance in each direction with its periodic images). Ions were added and concentrations were set to 1 mM MgCl_2_ and 150 mM KCl. The solvated and ionized system is composed of 781,332 atoms. The final conformation of the first set was selected for our current study. In accord with the study, all MD simulations were performed in NAMD using the CHARMM36 all-atom additive protein force field^35^ with a time step of 2 fs. Since simulations spanned 3 years, different versions of NAMD were used: 2.13^36^, 2.14, and 3^37^. 12 Å cut-off distance was used for van der Waals interactions. The particle-mesh Ewald method was used to calculate long-range electrostatic interactions. The temperature was kept constant at 310 K using a damping coefficient of 1 ps^−1^ for Langevin dynamics. The pressure was maintained at 1 atm using the Langevin Nosé–Hoover method with an oscillation period of 100 fs and a damping time scale of 50 fs.

Link3-4 of human dynein-2 in the pre-powerstroke conformation has an RMSD of 1.79 Å to the Link3-4 of the *D. discoideum* dynein-1 in the post-powerstroke conformation upon alignment of their C_α_ atoms. Superpositioning of Link1-2 of human dynein-2 in the pre-powerstroke conformation and *D. discoideum* dynein-1 in the post-powerstroke conformation shows minimal structural changes (RMSD of 1 Å), indicating that Link1-2 performs a rigid body movement during the priming stroke. The angle between the vector pointing from Link1-2 L1415 C_α_ atom to the L1312 C_α_ atom and the vector pointing from Link3-4 T1521 C_α_ atom to L1415 C_α_ atom were evaluated with four-quadrant inverse tangent and set as the linker angle, *θ*.

### SMD and cMD simulations

The starting system for the SMD simulations was selected from the equilibrated human dynein-2 motor domain conformations in the pre-powerstroke state. Constant velocity SMD simulations were performed, where a dummy atom binds with a virtual spring to the CoM of a group of ‘steered’ atoms (SMD atoms) and is pulled at a constant velocity (**v**) along the ‘pulling direction’ (**n**). During this pulling process, the following force is applied along the vector to the SMD atoms,^38^

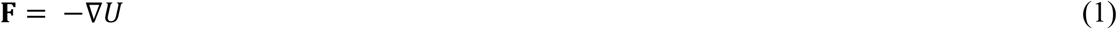

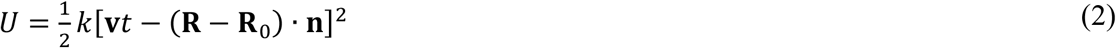

Here, *U* is the potential energy, *k* is the spring constant, *t* is time, and **R** and **R**_**0**_ are the coordinates of the center of mass of steered atoms at time *t* and 0, respectively.^38^ To define the SMD pulling vectors, dynein conformations were superimposed onto the ADP-bound post-power stroke structure via the Link 3-4 helices. C_α_ atoms of Link1-2 helices H4 (G1256-G1271), H5 (K1296-K1313), and H7 (E1322-W1349) were selected as SMD atoms. SMD vector was selected as the vector pointing from the CoM of the SMD atoms toward the CoM of the C_α_ atom coordinates of the H4, H5, and H7 helices of the ADP-bound structure (Fig. S10). The pulling velocity was set to 1 Å/ns and the spring constant was selected as 50 kcal/mol/Å^2^ in the SMD simulations, following the same parameters used for domain movement analysis in our recent study on the Spike protein.^39^ In these conditions, the CoM of SMD atoms closely followed the dummy atom, hence satisfying stiff-spring approximation,^40^ while the spring constant was soft enough to allow small deviations.

cMD simulations were performed after each SMD simulation to relieve excess stress and artificial deformations due to steering forces. Since Link-1-2 is expected to undergo rigid body motion (Supplementary Information),^15^ H4, H5, and H7 (steered atoms) of the starting conformation^22^ were superimposed on the end conformation of SMD simulation. Subsequently, the C_α_ atoms of the superimposed H4, H5, and H7 helices were selected as the first set of harmonic constraint centers to correct any deformation due to steering forces. Furthermore, a second set of harmonic constraints were applied to the C_α_ atoms of H4, H5, H7, H10 and H12 residues, which were kept fixed in the SMD simulations, at their SMD end conformation coordinates. A force constant of 1 kcal/mol/Å^2^ was used for each harmonic constraint.

### UMD Simulations and WHAM for Free-Energy Calculations

UMD simulations were performed using a modified potential energy function of the following form^29,30,41^

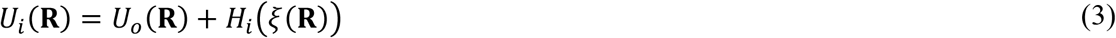

where **R** is the 3N-dimensional vector that defines protein structure, and N is the total atom number. *U*_*0*_(**R**) is the potential energy of the molecular system and 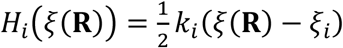 is a harmonic perturbing potential. The reaction coordinate ξ was defined as the RMSD of linker residues to their coordinates in the post-power stroke structure.^7^ To this aim, C_α_ atoms of Link3-4 H12, H13, S3, S4, S5, H14, and H15 (also used in the alignment of linker), and Link1-2 H4, H5, H7, and H10 were used for RMSD calculations and hence external bias was applied to these atoms. UMD simulations were initiated from end conformations of each SMD-cMD simulation cycle, and the equilibrium position of the applied harmonic constraint was selected as the ξ coordinate of the starting conformation. We also initiated simulations from infrequently sampled conformations of the first round. The harmonic restrain centers applied along the reaction coordinate are as follows: ξ = 3, 3.2, 3.6, 4.3, 4.9, 5.5, 6.1, 6.6, 7.2, 7.8, 8.4, 9.1, 9.7, 10.3, and 10.8 Å. A force constant of *k*_*i*_ = 25 kcal/mol/Å^2^ was used in all UMD simulations, except for 2 UMD simulations were centered at ξ =3.61 Å and 2.96 Å for which spring constants of 50 and 100 kcal/mol/Å^2^ were applied to sample these regions, respectively.

Using the distributions of the conformations sampled during UMD simulations by clustering them into Δξ=0.1 Å wide bins along the reaction coordinate, WHAM calculations were performed in WHAM v2.0.10^42^ to construct the free energy profile along the reaction coordinate with a convergence tolerance of 10^−6^ kcal/mol and histogram boundaries of 3 Å and 10.5 Å (See Supplementary Information). We calculated the average linker angles for all conformations within each WHAM RMSD bin and assigned these values to the corresponding free energy values of the bins.

### TMD simulations of an isolated linker

C_α_ atoms of linker residues were selected as the atoms to be guided via TMD.^31^ A bias potential was applied as a function of the instantaneous RMSD from the target coordinates with a spring constant of 200 kcal mol^-1^ Å^-2^. TMD simulation length was set to reach its target in 50 ns.

### Interaction analysis criteria

As was previously performed,^43,44^ a 6 Å cutoff distance between basic nitrogen and acidic oxygen was used for salt bridge formation,^45^ 8 Å cutoff distance between side chain carbon atoms for hydrophobic interactions,^46-48^ and a 3.5 Å cutoff distance and 30° cutoff angle for hydrogen bond formation,^49^ with pairs not meeting angle criteria of hydrogen bond formation classified as electrostatic interactions. Observation frequencies from MD simulations were classified as moderate (15-48%) or high (≥49%), with pairwise interactions below 15% excluded from further analysis.

## Author Contributions

Mert Gur (MG) and AY initiated the project. MG and Mert Golcuk (MGo) performed the molecular dynamics simulations. MG, MGo, SZY, and AY prepared the manuscript. MG supervised the project.

## Notes

The authors declare no competing financial interest.

## ACKNOWLEDGMENT

This work was supported by grants from the Partnership for Ad-vanced Computing in Europe (PRACE, PRA205144, MG), PRACE DECI-17 (Distributed European Computing Initiative) (17DECI0080), and Istanbul Technical University Scientific Re-search Projects (BAP) (MGA-2021-42803, MG), National Insti-tute of General Medical Sciences (GM094522 (AY), and the National Science Foundation (MCB-1617028 and MCB-1055017, AY). This work used resources, services, and support provided by Marconi100, MAHTI, and SAVIO.

## ABBREVIATIONS

ATP: Adenosine triphosphate
ADP: Adenosine di-phosphate
CC: coiled-coil
AAA: ATPases Associated with diverse cellular Activities
MT: Microtubule
MD: Molecular dynamics
SMD: steered molecular dynam-ics
TMD: Targeted molecular dynamics
UMD: Um-brella sampling molecular dynamics

