## SupplementaryMaterial for "The Mechanism and Energetics of the Dynein Priming Stroke"

### Structural Mechanism and Energetics of the Cytoplasmic Dynein Priming Stroke

[*]Corresponding Author

#### Supplementary Methods

##### Weighted Histogram Analysis Method (WHAM) and Error Analysis

Upon performing umbrella sampling molecular dynamics (UMD) simulations with modified potential energy functions, a set of biased probability distributions $P_i^{(b)}(\xi)$ can be obtained as

$$P_i^{(b)}(\xi) = exp[-\beta\Phi_i(\xi)] = \langle\delta[\xi(\mathbf{R}) - \xi_i]\rangle \qquad (1)$$

where $\Phi_i(\xi)$ is the free energy at reaction coordinate ξ when the simulation is carried out with the perturbing potential $H_i(\xi(\boldsymbol{R}))$. From MD trajectories, $P_i^{(b)}(\xi)$ is computed as the normalized histogram of the values occurring during simulation *i*. The corresponding unbiased probability distribution is defined as,

$$P_i^{(u)}(\xi) = exp[\beta(H_i(\xi) - f_i)]P_i^{(b)}(\xi) \qquad (2)$$

where $f_i$ is the free energy coming from the adding of the biasing potential $H_i(\xi(\mathbf{R}))$ to $U_o(\mathbf{R})$. The WHAM[1] was used to recombine the unbiased histograms $P_i^{(u)}(\xi)$ obtained from UMD simulations to obtain the total probability distribution $P_o(\xi)$:[2]

$$P_o(\xi) = C\sum_{i=1}^{N}\frac{n_i}{\sum_{j=1}^{N} n_j exp[-\beta(H_j(\xi) - f_j)]}P_i^{(b)}(\xi) \qquad (3)$$

where *C* is a normalization constant, *N* is the total number of MD simulations, and $n_i$ is the number of coordinate sets in the $i^{th}$ MD simulation to compute $P_i^{(b)}(\xi)$. All terms except the free energy parameters $\{f_i\}$ can be directly computed. To evaluate $\{f_i\}$, the following equation is solved iteratively,

$$exp[-\beta f_i] = C\int d\xi\sum_{i=1}^{N}\frac{n_i exp[-\beta H_k(\xi)]}{\sum_{j=1}^{N} n_j exp[-\beta(H_j(\xi) - f_j)]}P_i^{(b)}(\xi) \qquad (4)$$

From $P_o(\xi)$, the free energy surface along ξ can be evaluated by $\Phi_i(\xi) = -\frac{1}{\beta}ln[P_o(\xi)]$ .

Error analysis were performed using the bootstrap method. 15 “trajectories” were generated by randomly selecting 100,000 data points from our UMD simulations (setting correlation time to 0.5 ns) and constructing the free energy surfaces. The standard deviation of these trajectories was in the range 0.01-0.2 kcal/mol for each bin along the reaction coordinate, hence indicating low error in our WHAM calculations.

**Principal Components Analysis (PCA) and Free Energy Landscape Calculations.** Linker conformations sampled during MD simulations were aligned with the pre-powerstroke conformation of human dynein-2 using the $C_\alpha$ atoms of Link3-4 helices and beta sheets. Using the aligned linker coordinates, the covariance matrix is generated as follows,

$$\mathbf{C} = \langle(\mathbf{R} - \langle\mathbf{R}\rangle)(\mathbf{R} - \langle\mathbf{R}\rangle)^{\mathrm{T}}\rangle \tag{5}$$

where $\mathbf{R}$ is the $3 \times 407 = 1221$ dimensional configurational vector composed of the instantaneous $C_\alpha$ atom coordinates of the linker and $\langle\mathbf{R}\rangle$ is the trajectory average of **R**. Eigenvalue decomposition of **C** is performed and principal components (PCs) are obtained as follows,

$$\mathbf{C} = \sum_{i=1}^{3N} \sigma_i \mathbf{p}_i \, \mathbf{p}_i^{T} \tag{6}$$

Here, $\mathbf{p}_i$ is the $i^{\text{th}}$ PC and $\sigma_i$ is the the corresponding variance. Thus, $\sigma_i$ scales with the magnitude of motion along $\mathbf{p}_i$ . PCs are ordered in descending order with respect to their $\sigma_i$ values. Since PC1 and PC2 have the largest variances, $\mathbf{p}_1$ and $\mathbf{p}_2$ are the most dominant motions observed in the MD simulations. Projections of instantaneous coordinates along PCs are evaluated as $\mathbf{r}_i = \mathbf{p}_i^{T}(\mathbf{R} - \langle\mathbf{R}\rangle)$.

Distributions of the projections $f(\mathbf{R})$ were computed along PC1 and PC2. Higher positive values indicate increased levels of linker becoming a straight conformation, whereas lower negative values represent increased degree of linker bending. Using distributions along PC1 and PC2, free energy surfaces were calculated as, $A(\mathbf{R}) = -kTln\left(f(\mathbf{R})\right) + \text{constant}$.[3]

#### Supplementary Tables

**Table S1. Observation frequency of intradomain interactions of the linker along ξ.** Interactions that take place between Link1-2 and Link3-4 are shown in black, while interactions between the hinge and the remaining linker are showed in red. Those interactions occurring exclusively in the semi-bent, transition state and, and bent conformations of the linker are highlighted in cyan, magenta, and brown, respectively. The darkness level of the blue color represents the frequency of these interactions depending on the root mean square deviation (RMSD) (left panels) and the angle (right panels) of the linker.

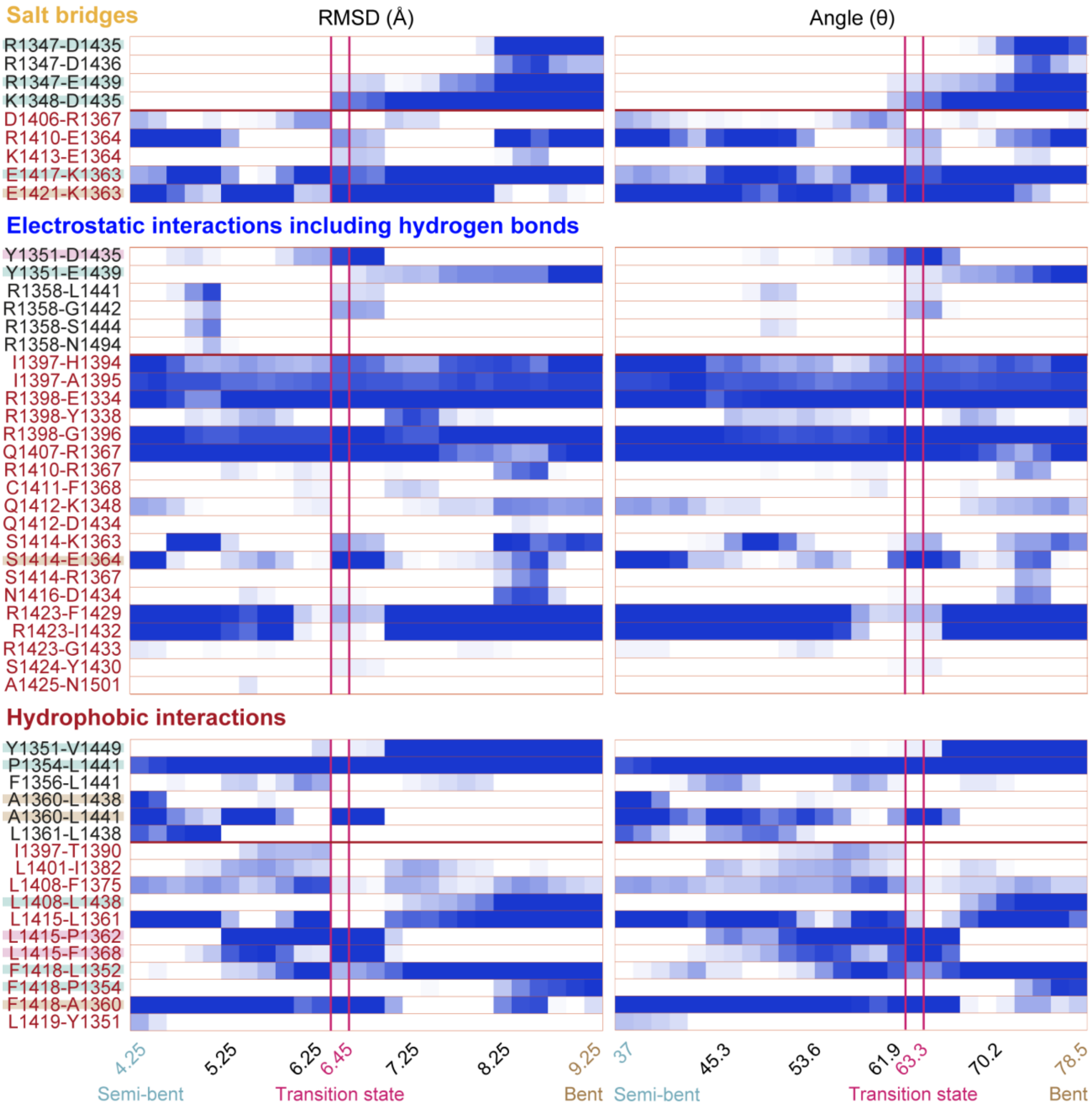

**Table S2. Observation frequency of interdomain salt bridge interactions between the linker and the AAA+ ring along ξ.** Interactions occurring between linker and AAA1-6, are shown in dark blue, light blue, green, yellow, orange, and pink, respectively. Those interactions occurring exclusively in the semi-bent, transition state and, and bent conformations of the linker are highlighted in cyan, magenta, and brown, respectively. The darkness level of the blue color represents the frequency of these interactions depending on the RMSD (left panels) and the angle (right panels) of the linker.

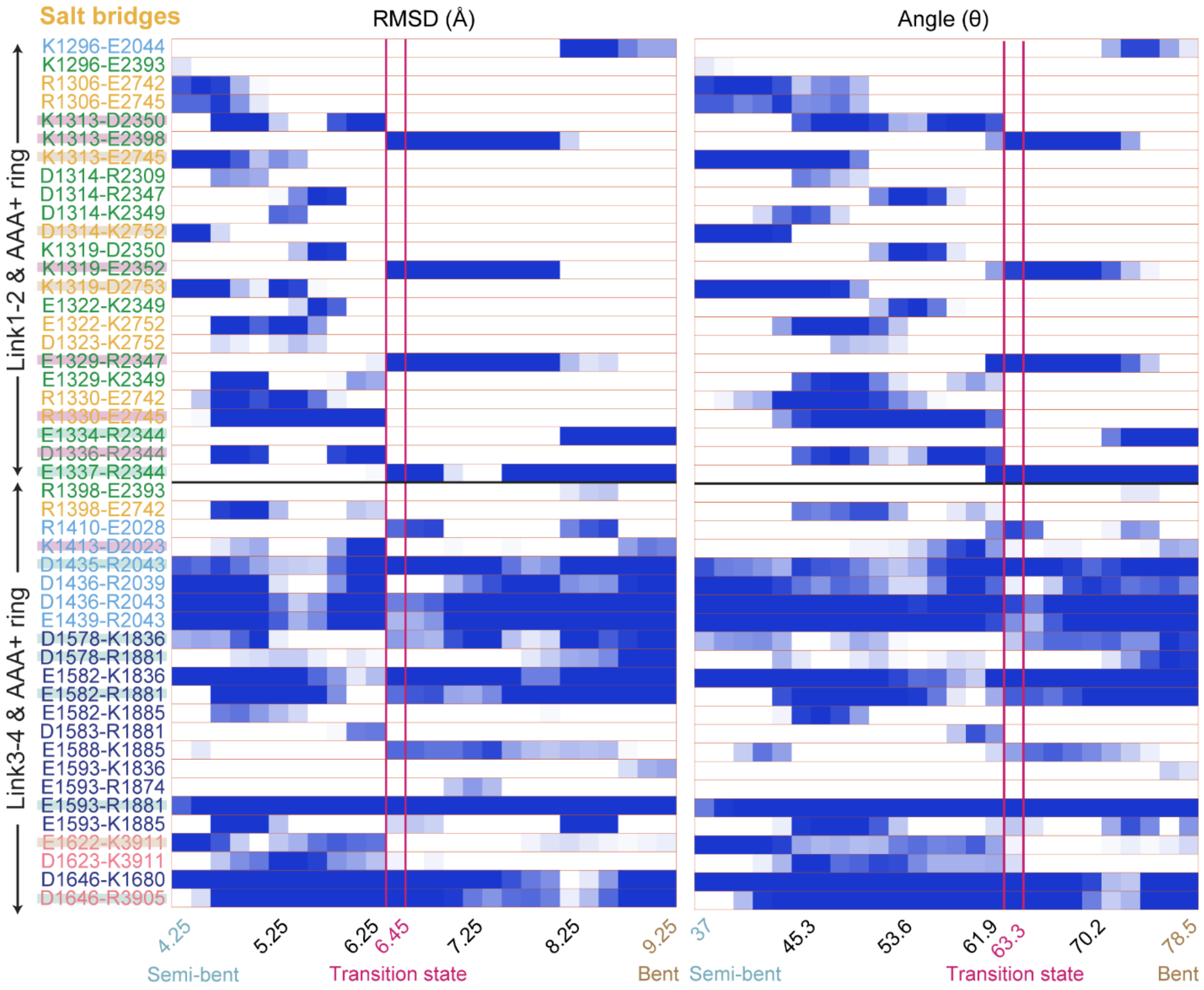

**Table S3. Observation frequency of interdomain electrostatic interactions (including hydrogen bonds) between the linker and the AAA+ ring along ξ.** Interactions occurring between linker and AAA1-6, are shown in dark blue, light blue, green, yellow, orange, and pink, respectively. Those interactions occurring exclusively in the semi-bent, transition state and, and bent conformations of the linker are highlighted in cyan, magenta, and brown, respectively. The darkness level of the blue color represents the frequency of these interactions depending on the RMSD (left panels) and the angle (right panels) of the linker.

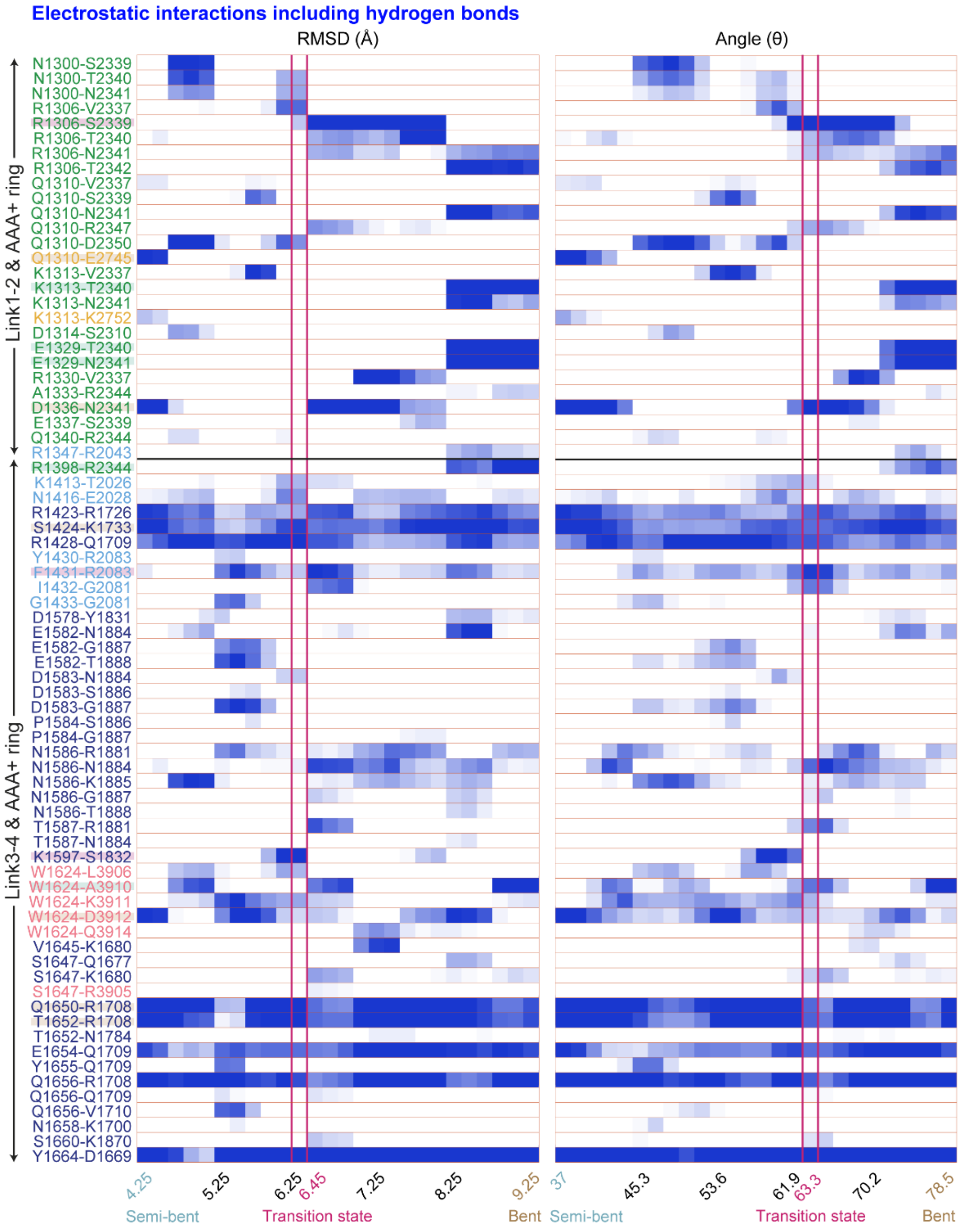


**Table S4. Observation frequency of interdomain hydrophobic interactions between the linker and the AAA+ ring along ξ.** Interactions occurring between linker and AAA1-6, are shown in dark blue, light blue, green, yellow, orange, and pink, respectively. Those interactions occurring exclusively in the semi-bent, transition state and, and bent conformations of the linker are highlighted in cyan, magenta, and brown, respectively. The darkness level of the blue color represents the frequency of these interactions depending on the RMSD (left panels) and the angle (right panels) of the linker.

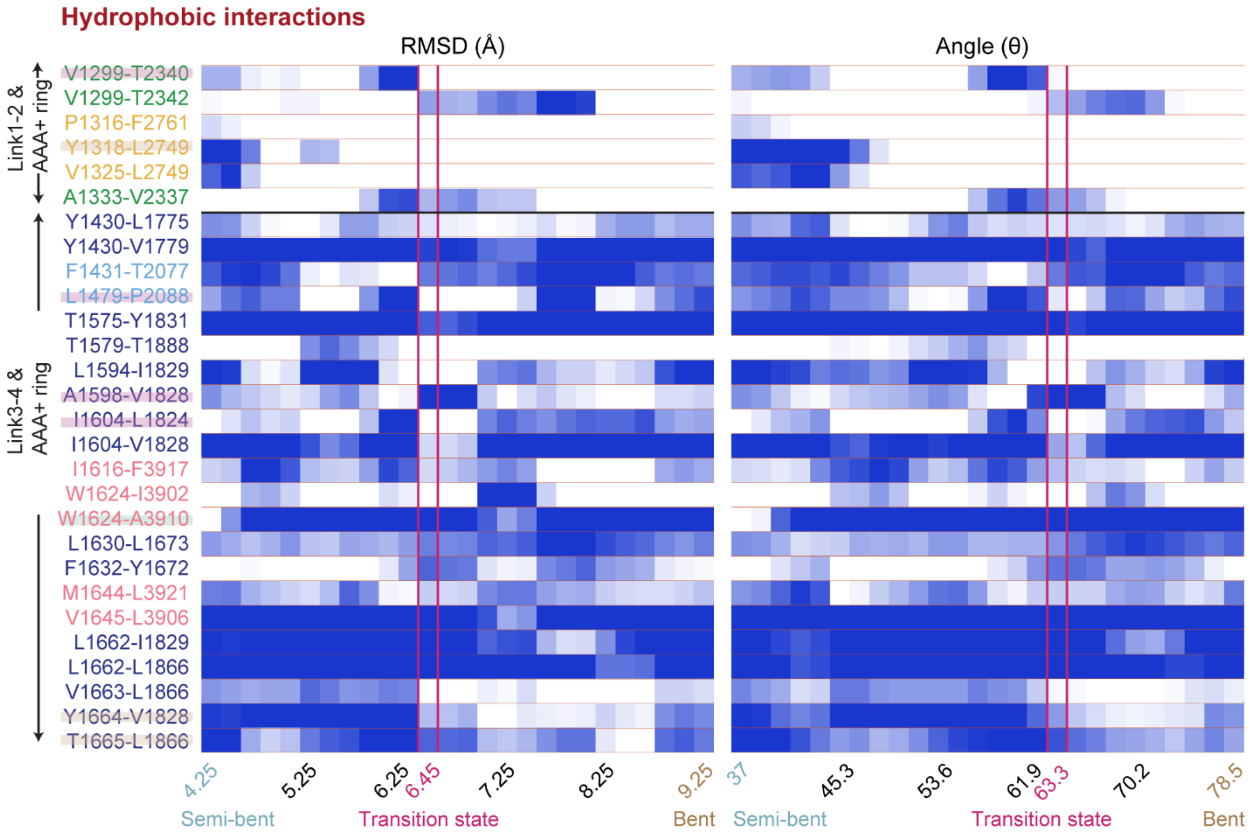

**Table S5. Observation frequency of intradomain interactions for isolated linker between the linker and the AAA+ ring along ξ.** Interactions that take place between Link1-2 and Link3-4 are shown in black, while interactions between the hinge and the remaining linker are showed in red. Those interactions occurring exclusively in the semi-bent, transition state and, and bent conformations of the linker are highlighted in cyan, magenta, and brown, respectively. The darkness level of the blue color represents the frequency of these interactions depending on the RMSD (left panels) and the angle (right panels) of the linker.

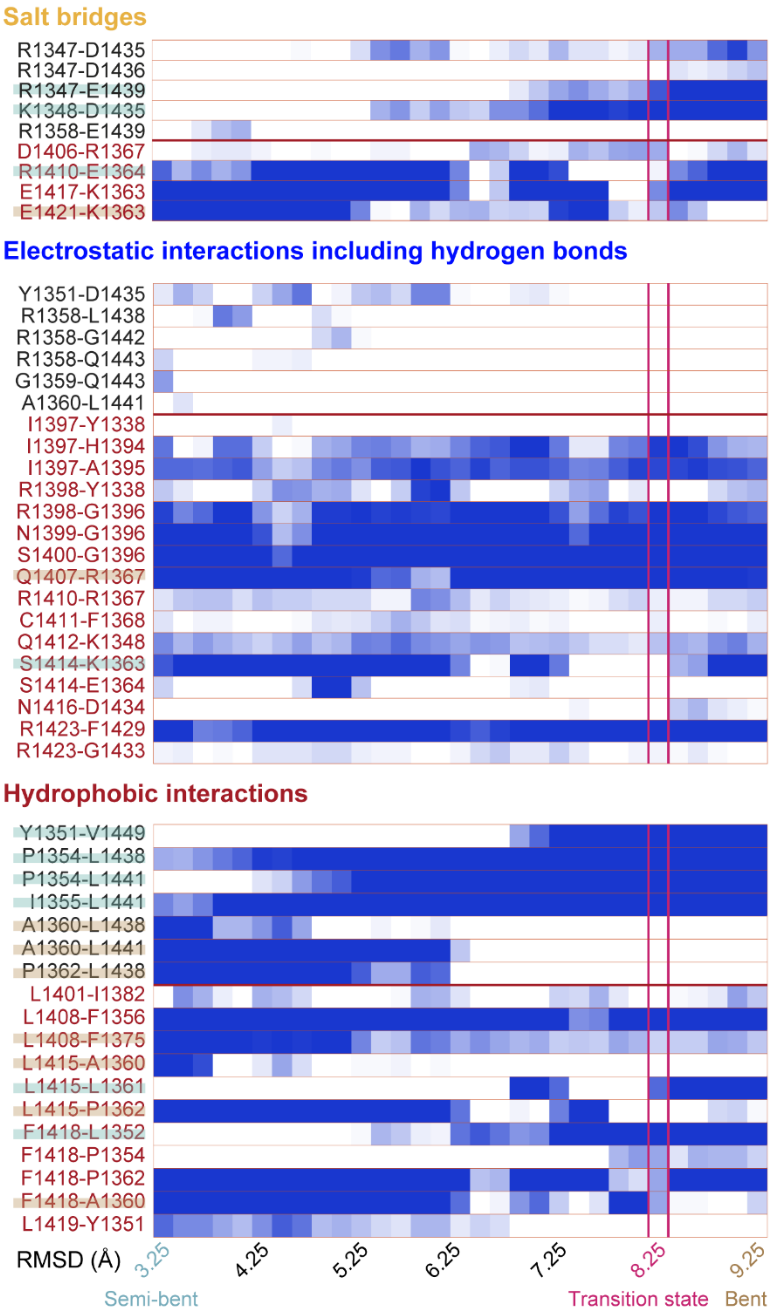


#### Supplementary Figures

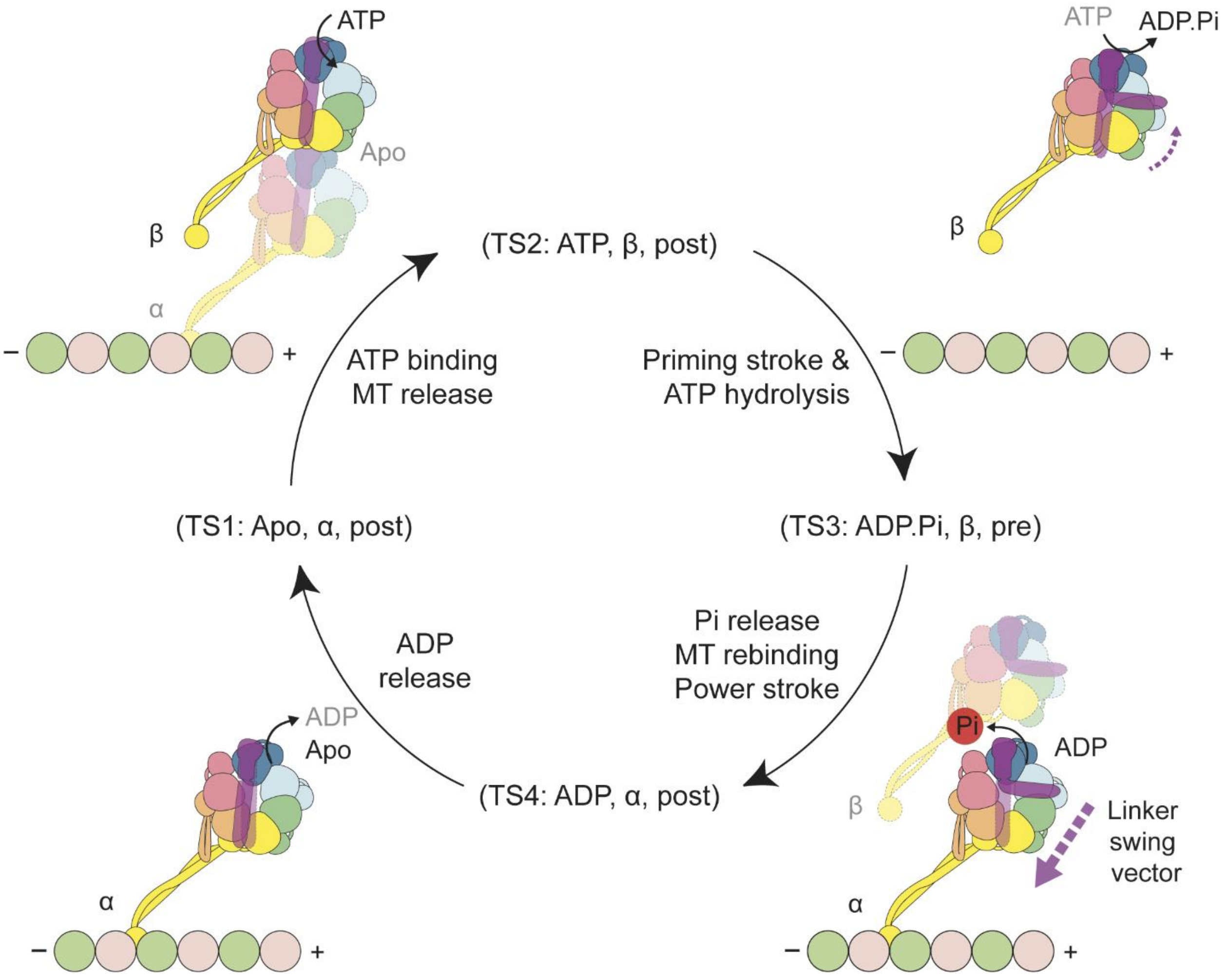


**Fig. S1. Mechanochemical cycle of the dynein motor domain.**

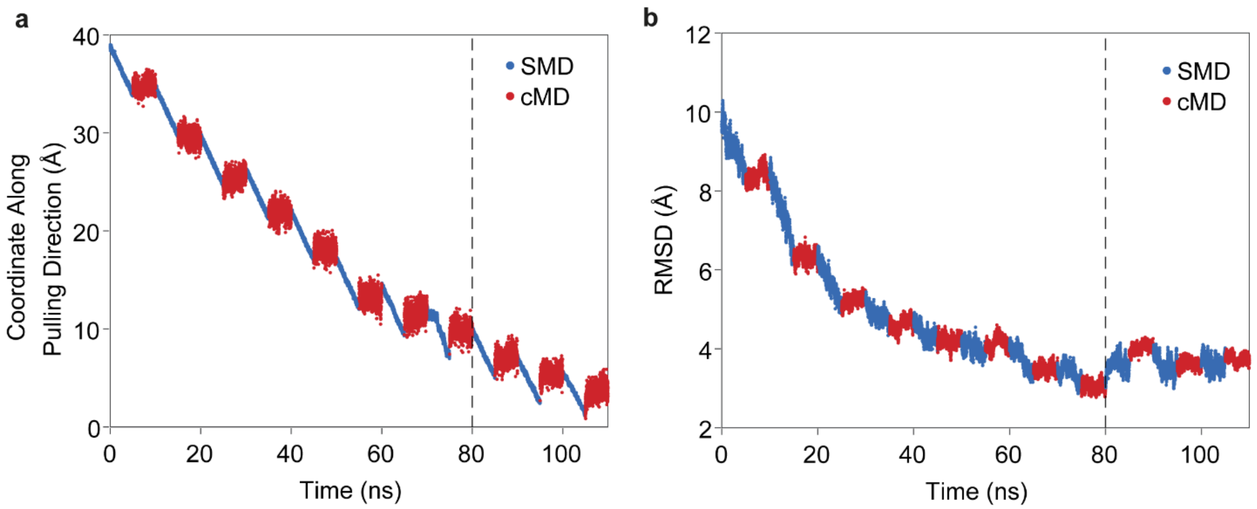


**Fig. S2. Progress of steered MD (SMD) simulation protocol towards the target linker structure.** **a,** The distance between the center of mass (CoM) of SMD atoms to their straight linker structure[4] in SMD and constrained MD (cMD) simulations. **b,** RMSD between linker conformations produced during SMD-cMD simulations and the straight structure of the linker. H4, H5, H7, H10, H12, H13, S3, S4, S5, H14, and H15 $C_\alpha$ atoms were used for RMSD calculations. Conformations sampled in SMD simulation were aligned to the straight structure via Link3-4 H12, H13, S3, S4, S5, H14, and H15 $C_\alpha$ atoms. The SMD-cMD cycles did not progress towards their target any further after 80 ns (the dashed vertical line).

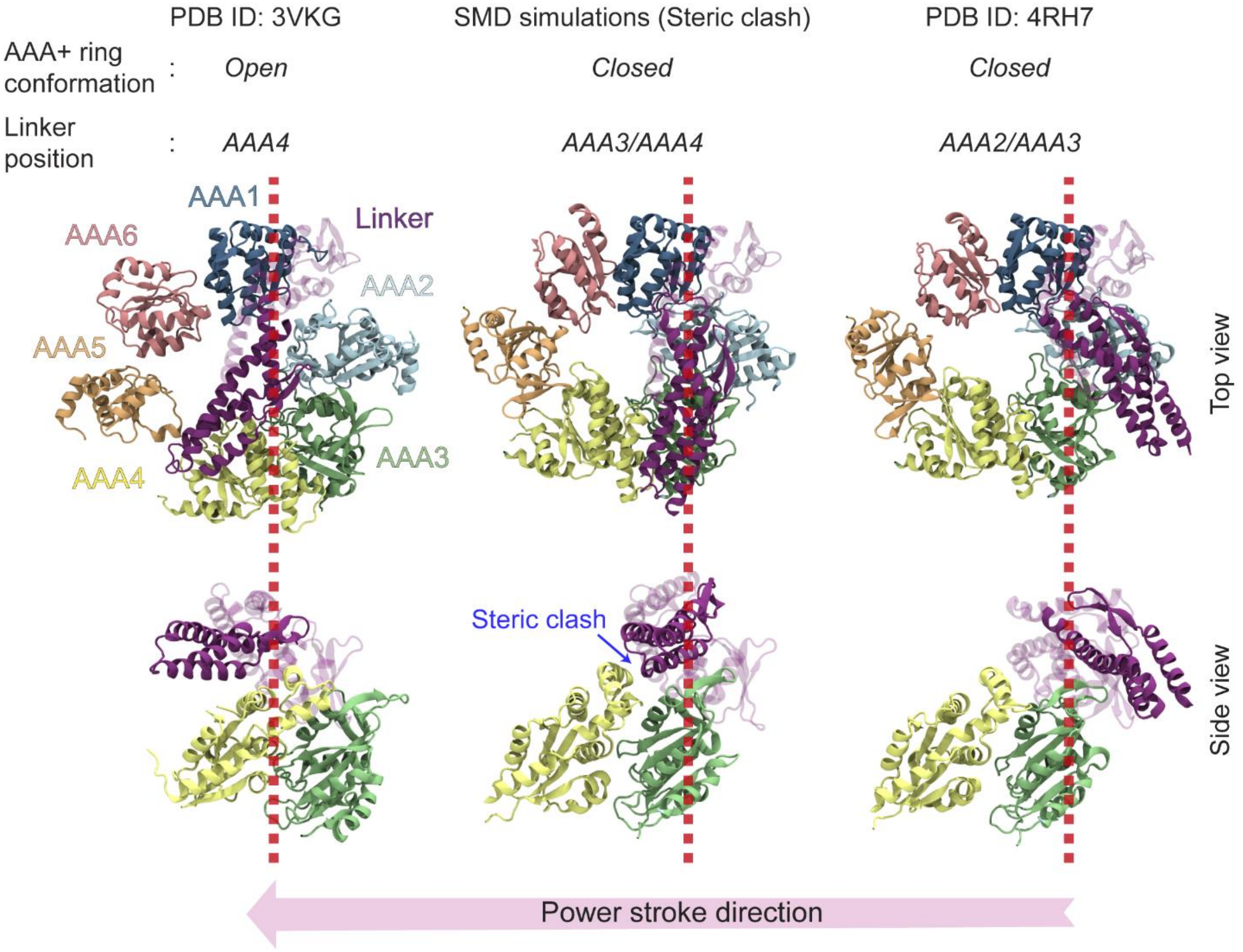


**Fig. S3. SMD simulations in the direction of the powerstroke of the linker.** (Top) The AAA+ ring and linker conformations in the post-powerstroke state of *Dictyostelium discoideum* dynein-1 (PDB ID: 3VKG[4]), the final conformation of SMD simulations, and pre-powerstroke conformation of human dynein-2 (PDB ID: 4RH7[5]). (Bottom) The side view of Link1-2, AAA3 and AAA4 shows that the SMD were not able to progress to the straight conformation of the linker due to a steric clash (blue arrow) with AAA4. The red dashed line that passes through the hinge region of the linker highlights the structural transformations of Link1-2 relative to Link3-4.

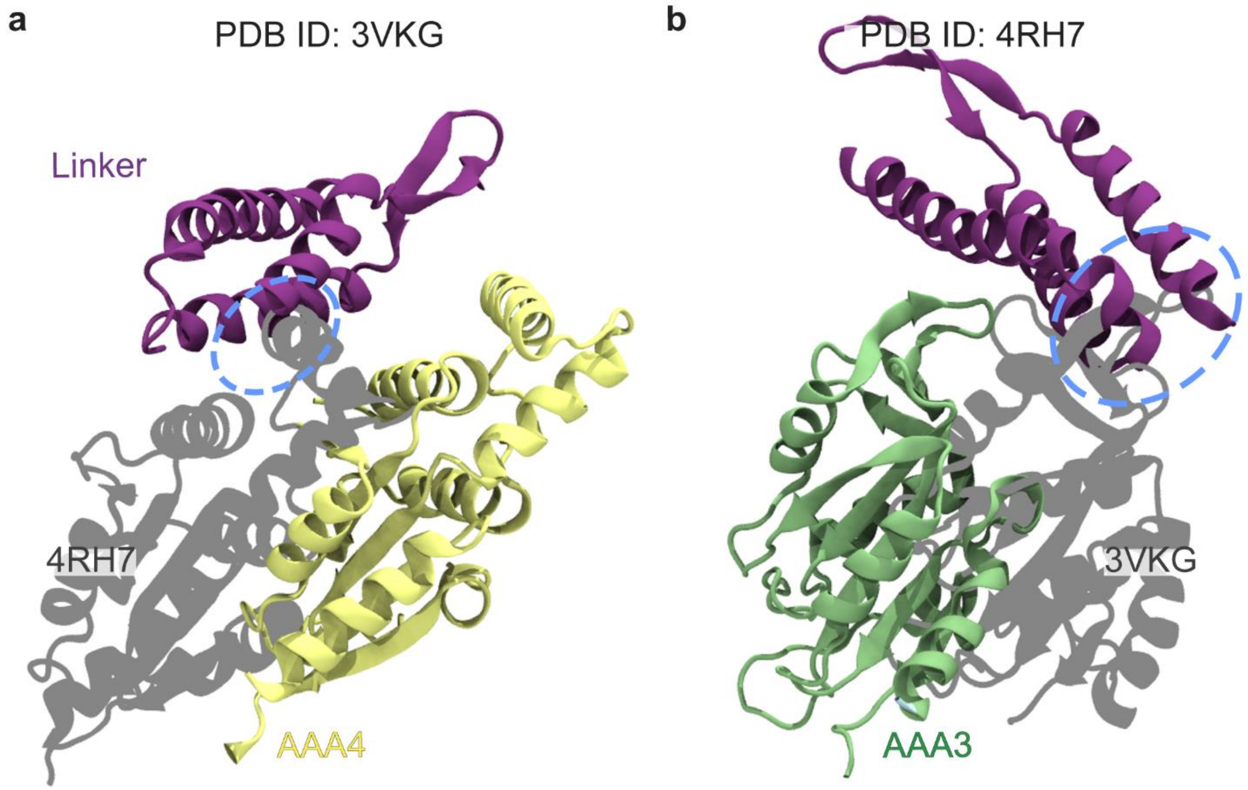


**Fig. S4. Steric clash between the linker and AAA domains upon superpositioning of the pre- and post-powerstroke conformations of the dynein motor domain.** Structures of *D. discoideum* dynein-1 in the ADP-bound state (PDB ID: 3VKG[4]) and human dynein-2 in the ADP.Pi state (PDB ID: 4RH7[5]) were superimposed with respect to Link3-4, as it was previously shown by Schmidt et al.[5]. **a,** The movement of the linker in the powerstroke direction on the surface of the ADP.Pi state of the AAA+ ring results in a steric clash with the AAA4 site. **b,** The movement of the linker in the priming stroke direction on the surface of the ADP bound conformation of the AAA+ ring results in a steric clash with the AAA3 site.

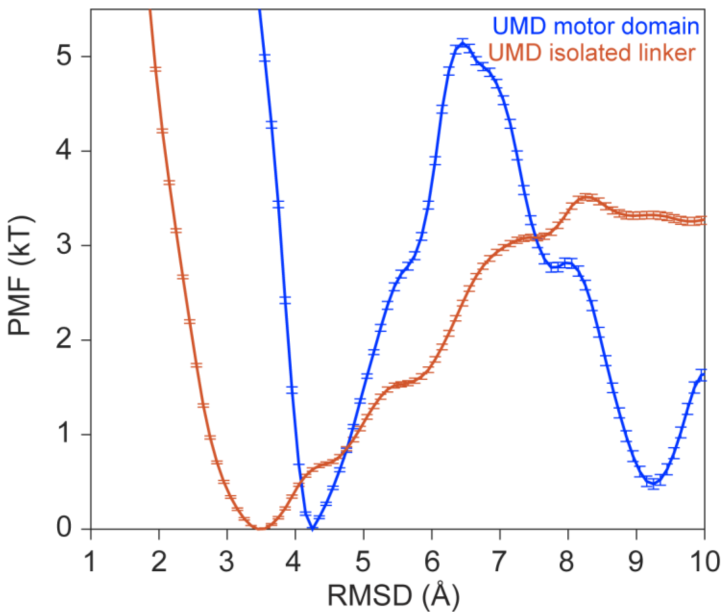


**Fig. S5. Error bars for the free energy surface generated using UMD simulations of the full-length dynein motor domain and isolated linker.** Error bars obtained via bootstrap analysis are shown on the free energy profiles along the reaction coordinate of the linker priming movement.

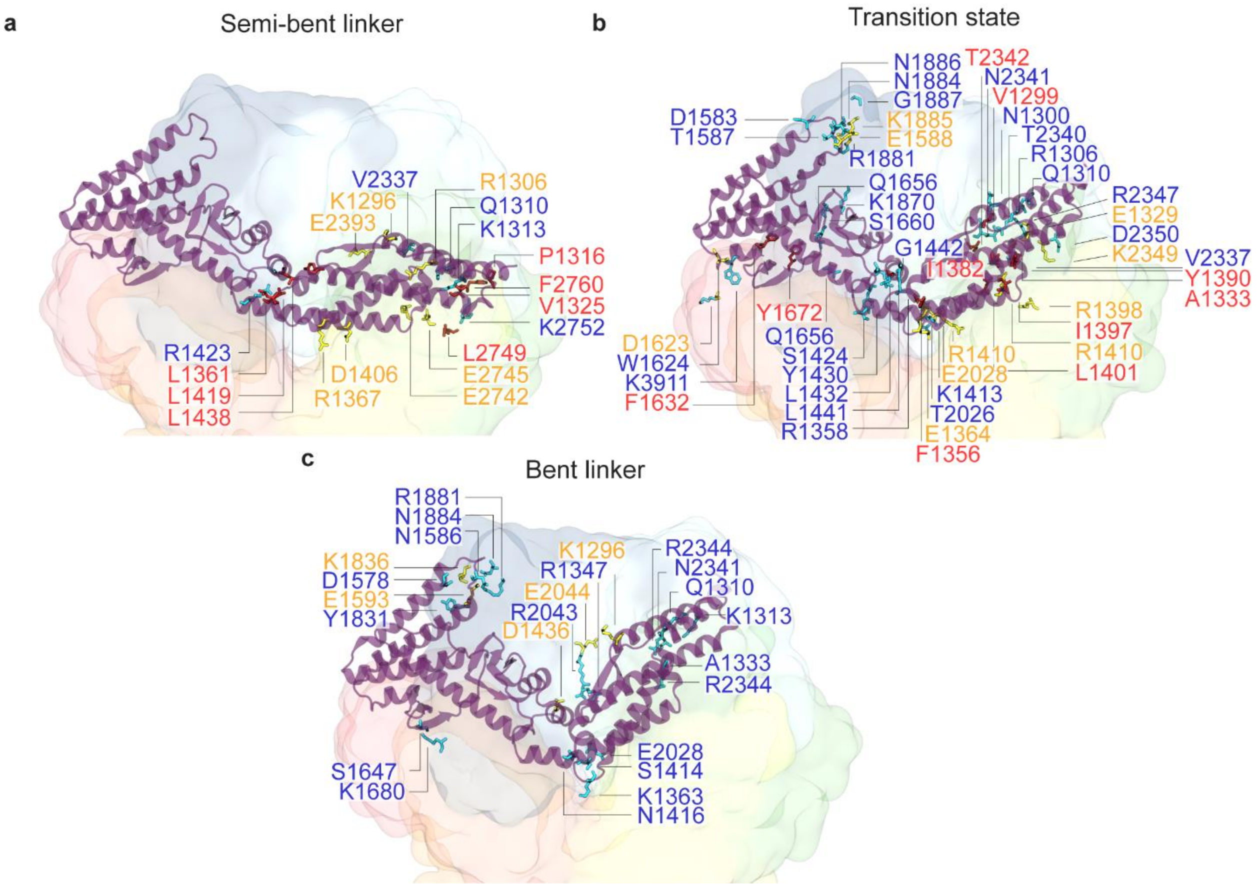


**Fig. S6. The medium frequency intradomain interactions of the linker and interdomain interactions between the linker and the AAA+ ring.** Location of forming and breaking medium frequency interactions are shown for the semi-bent **(a),** transition state **(b),** and bent **(c)** conformations of the linker. Hydrophobic interaction pairs are denoted by red, electrostatic interaction pairs by blue, and salt bridge interaction pairs by yellow.

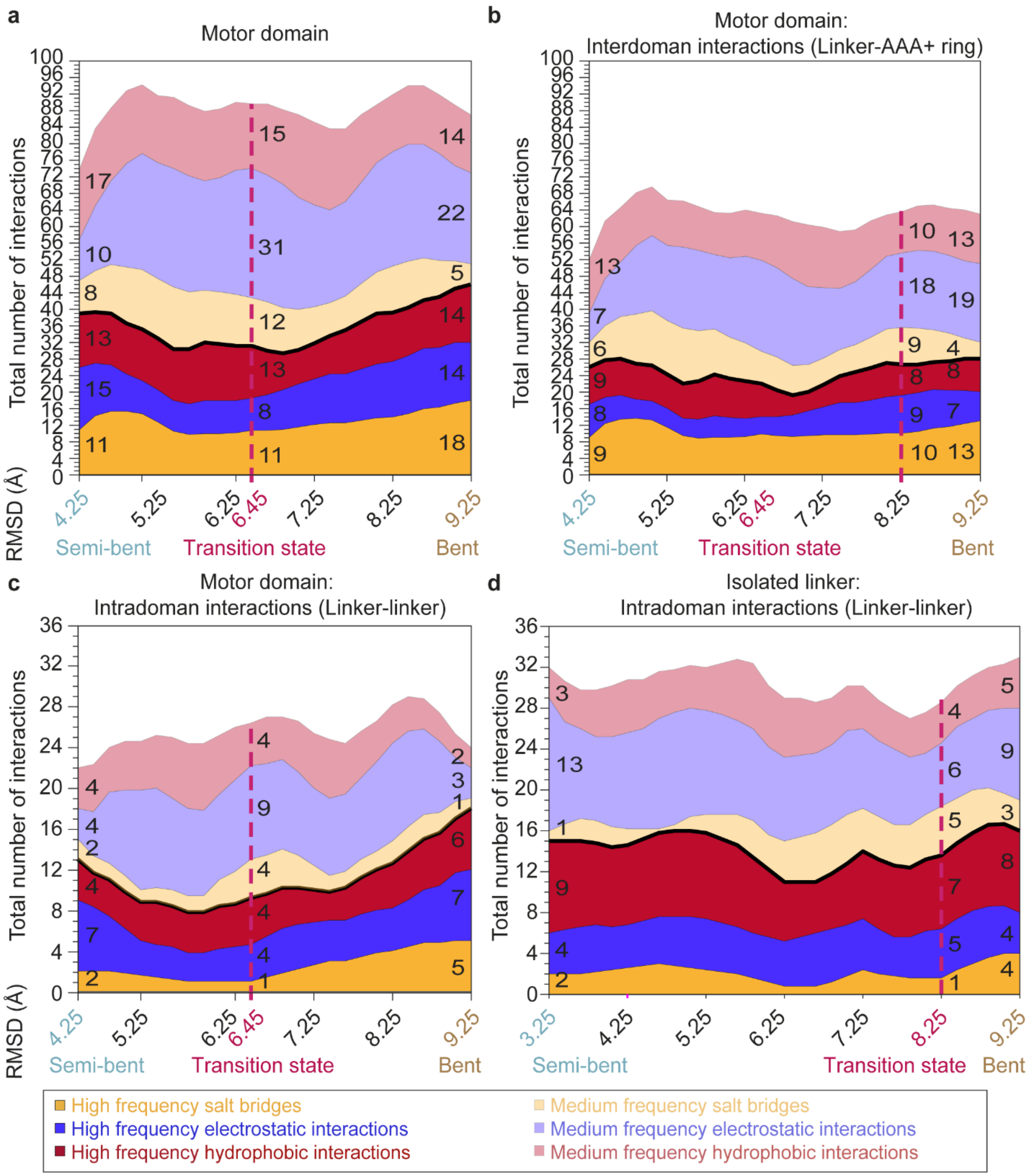


**Fig. S7. Change in the total number of intradomain and interdomain interactions of the linker during its priming movement. a-d,** Changes in the total number of salt bridges, electrostatic interactions (including hydrogen bonds), and hydrophobic interactions observed with high frequency (darker color) and medium frequency (lighter color), due to the formation and breakage of these interactions during the priming movement of the linker. The numbers of interactions are shown for each type of interactions.

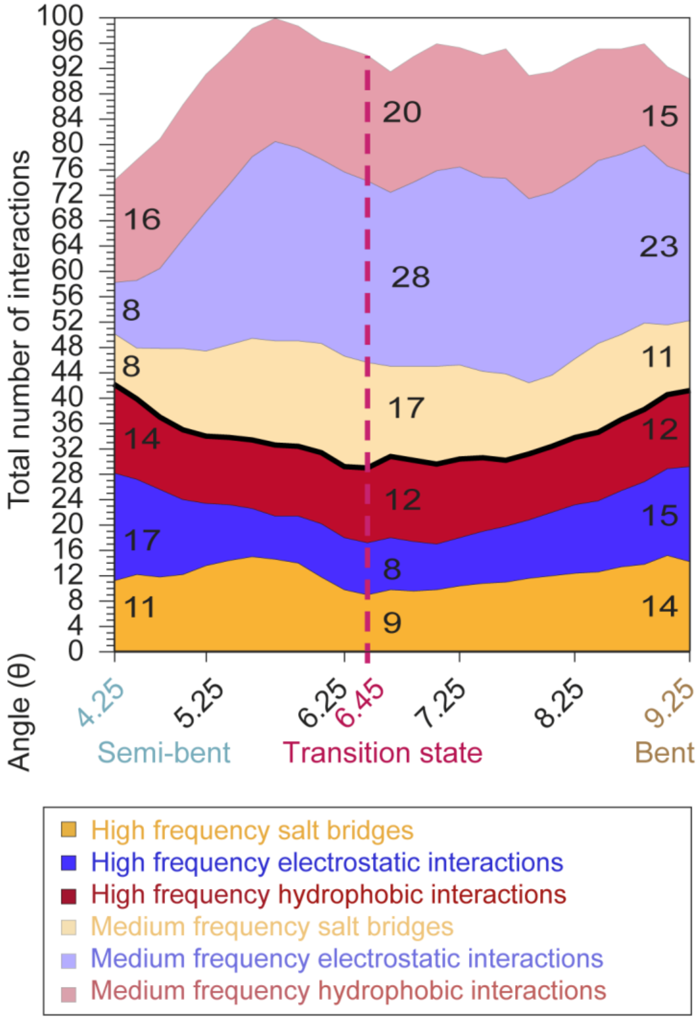


**Fig. S8. The changes in the number of pairwise interactions based on the linker angle.** Changes in the total number of salt bridges, electrostatic interactions (including hydrogen bonds), and hydrophobic interactions observed with high frequency and medium frequency are shown with darker and lighter colors, respectively.

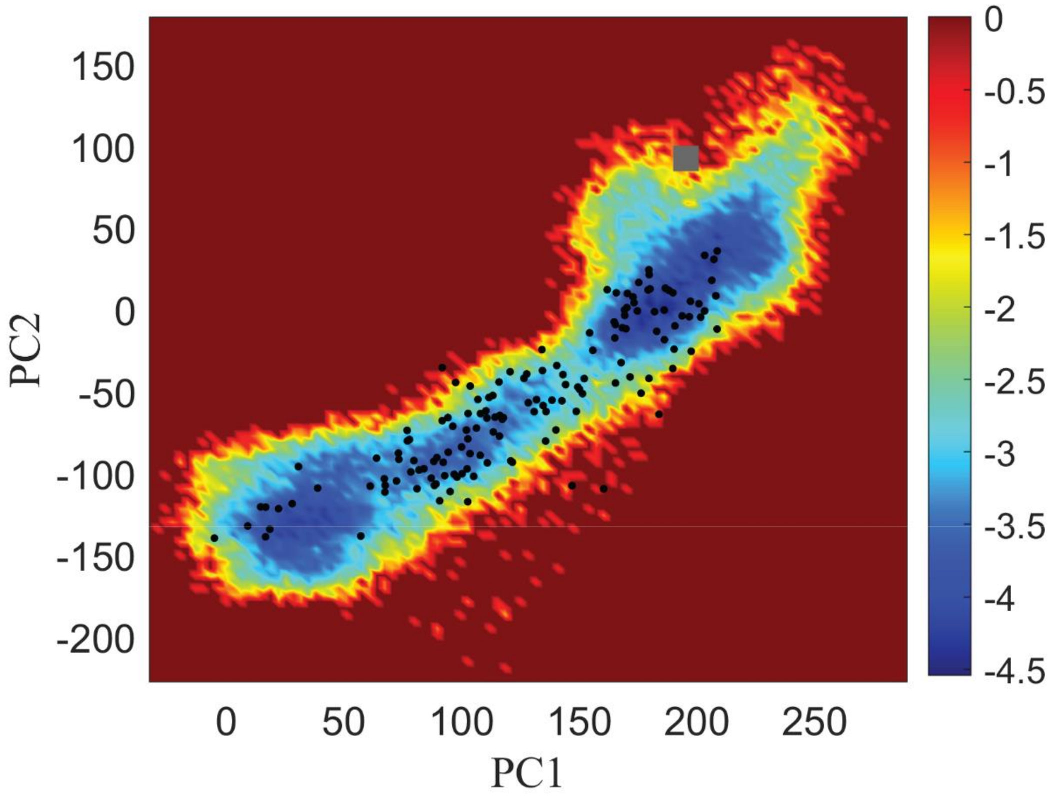


**Fig. S9. Trajectory of unconstrained MD simulations initiated from the close proximity of the energy barrier.** 21 unconstrained MD simulations were initiated from the close proximity of the energy barrier. The black dots show the conformations sampled in these simulations, with each conformation separated by 25 ns in time

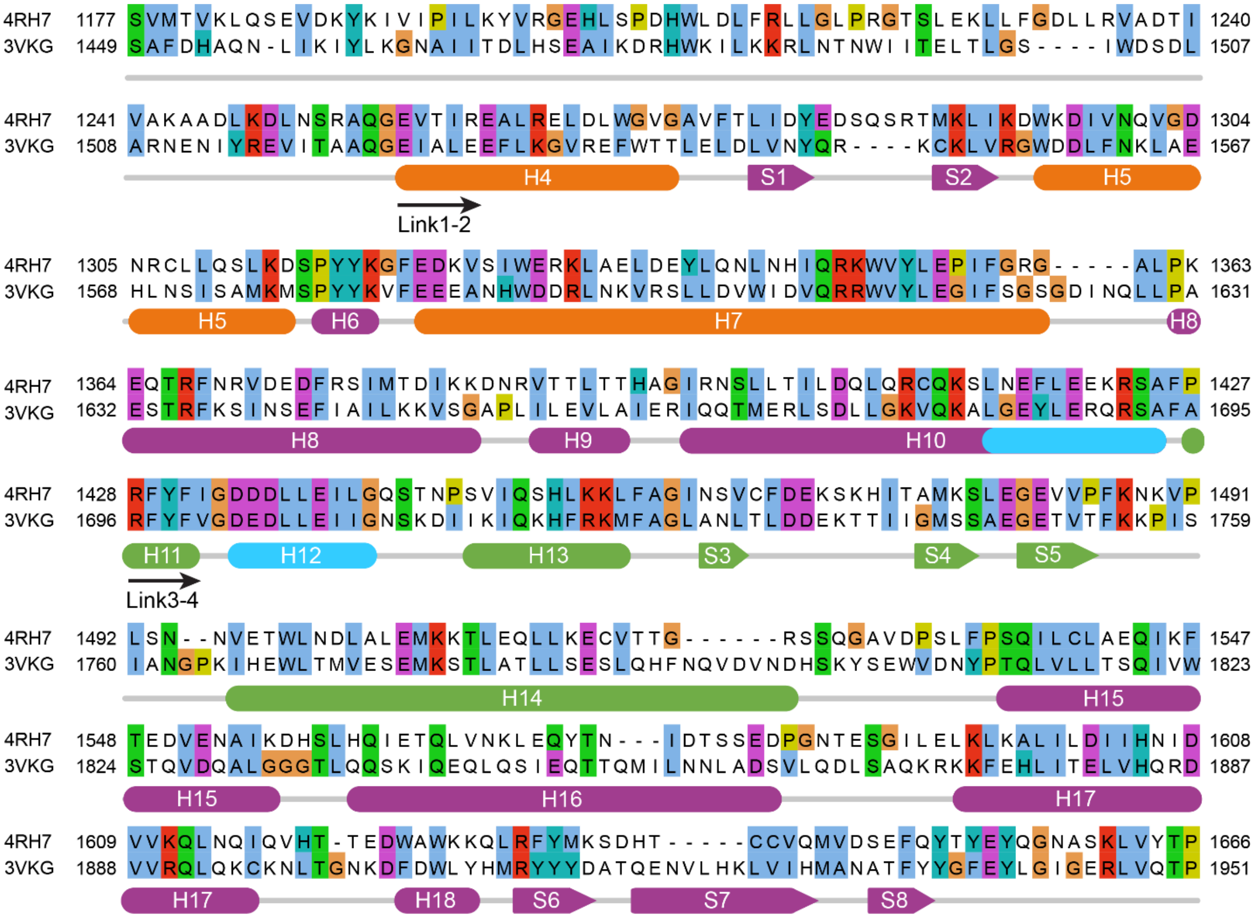

**Fig. S10. Sequence alignment of dynein-2 bent linker and dynein-1 straight linker.** Sequence alignment of human cytoplasmic dynein-2 and *D. discoideum* cytoplasmic dynein-1 linkers. Location of alpha helices (H4-H18) and beta sheets (S1-S8) are indicated on the alignment.
